# NIMBus: a Negative Binomial Regression based Integrative Method for Mutation Burden Analysis

**DOI:** 10.1101/2020.05.29.124149

**Authors:** Jing Zhang, Jason Liu, Patrick McGillivray, Caroline Yi, Lucas Lochovsky, Donghoon Lee, Mark Gerstein

## Abstract

**Background:** Identifying frequently mutated regions is a key approach to discover DNA elements influencing cancer progression. However, it is challenging to identify these burdened regions due to mutation rate heterogeneity across the genome and across different individuals. Moreover, it is known that this heterogeneity partially stems from genomic confounding factors, such as replication timing and chromatin organization. The increasing availability of cancer whole genome sequences and functional genomics data from the Encyclopedia of DNA Elements (ENCODE) may help address these issues.

**Results:** We developed a Negative binomial regression-based Integrative Method for mutation Burden analysiS (NIMBus). Our approach addresses the over-dispersion of mutation count statistics by (1) using a Gamma-Poisson mixture model to capture the mutation-rate heterogeneity across different individuals and (2) estimating regional background mutation rates by regressing the varying local mutation counts against genomic features extracted from ENCODE.

We applied NIMBus to whole-genome cancer sequences from the PanCancer Analysis of Whole Genomes project (PCAWG) and other cohorts. It successfully identified well-known coding and noncoding drivers, such as TP53 and the TERT promoter. To further characterize the burdening of non-coding regions, we used NIMBus to screen transcription factor binding sites in promoter regions that intersect DNase I hypersensitive sites (DHSs). This analysis identified mutational hotspots that potentially disrupt gene regulatory networks in cancer. We also compare this method to other mutation burden analysis methods.

**Conclusion:** NIMBus is a powerful tool to identify mutational hotspots. The NIMBus software and results are available as an online resource at github.gersteinlab.org/nimbus.

## Background

Population-level analysis, which looks for regions mutated more frequently than expected, is one of the most powerful ways of identifying deleterious mutations in diseases [1-3]. The availability of whole genome sequencing (WGS) has provided unprecedented statistical power to perform such analyses. An accurate quantification of mutation burden is important to help uncover the genetic cause of various diseases, which in turn would allow for targeted therapies in clinical studies. One typical application of such analysis is to find burdened regions in cancer genomes as potential drivers.

However, mutation burden tests for somatic variants in cancer research remain challenging for several reasons. First, it is well known that cancer genomes are heterogeneous [4]. If a constant mutation rate is assumed, the positional level mutation counts often demonstrate larger than expected variance, known as overdispersion. This assumption results in poor data fitting and can lead to numerous false positives [5], so it is necessary to introduce more sophisticated models to handle this mutation rate heterogeneity. Second, numerous genomic features have been reported to largely affect the mutation process [6-12], necessitating careful correction in burden analysis. These features include chromatin status and replication timing. Various strategies have been suggested for integrating these features to calibrate background mutation rate [7, 13-18]. However, these strategies may be limited by both the number and kinds of features used to model background mutation rate. For instance, cancer cells are usually highly heterogeneous and thus are not necessarily matched by features from a single cell type. Moreover, assay data does not necessarily exist for each feature type in all cell types. Lastly, many studies have shown that noncoding mutations can serve as driver events for diseases. For example the mutations in the TERT promoter were found to be associated with cancer progression [19-21]. A recent study of non-coding mutations in breast cancer identified driver mutations in three genes – FOXA1, NEAT1, and RMRP [22]. Hence, unified analysis of coding and noncoding regions is needed to give a thorough annotation of discovered hotspots.

Here, we propose a Negative binomial regression based Integrative Method for mutation Burden analysiS (NIMBus) that addresses the three problems mentioned above. It first intuitively treats mutation rates from different individuals as random variables with a gamma distribution, and resultantly models the pooled mutation counts from a heterogeneous population as a negative binomial distribution to handle overdispersion. Furthermore, to capture the effect of covariates, NIMBus integrates extensive features in all available tissues from Roadmap Epigenomics Mapping Consortium (REMC) and the Encyclopedia of DNA Elements (ENCODE) project to create a covariate matrix to predict the local mutation rate with high precision through regression.

In addition, NIMBus was used to analyze the most comprehensive noncoding annotations from ENCODE in two ways. First, our approach enabled us to effectively pinpoint mutation hotspots associated with disease progression and to better understand the associated biological mechanisms. This was accomplished by applying our method to the transcription start site (TSS) regions. Second, NIMBus targeted key transcription factor binding sites to give insight into the potential mechanisms for transcriptional regulation. Lastly, we compared our results to those from ICGC/TCGA Pan-Cancer Analysis of Whole Genomes Consortium [23]. To better illustrate how NIMBus works, Figure 1 gives its workflow.

**Figure 1.**
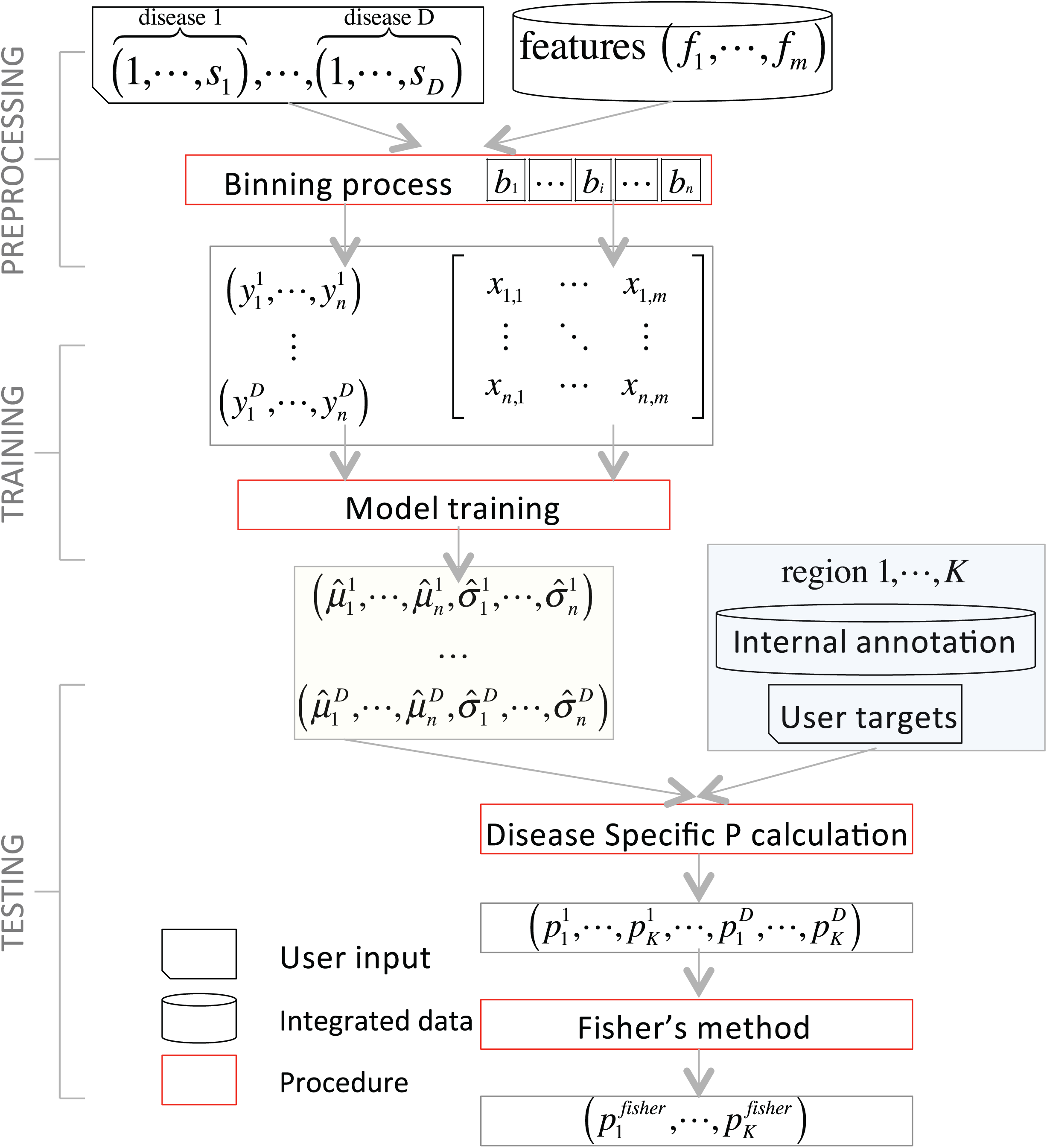
Flowchart of NIMBus. For a given disease *d* (1 ≤ *d* ≤ *D*), *s*_*d*_ represents the total number of samples for that disease. In addition, there are a total of m features which are denoted as *f*_l_ … *f*_m_. The mutations from the samples and the features are binned on the bins *b*_l_ … *b*_n_ for a total of n. Two resulting matrices are produced, ***Y*** and ***X***. The matrix ***Y*** is a *D* x *n* matrix consisting of mutation counts while ***X*** is an n × m matrix consisting of feature values. Training the negative binomial model gives, for each disease, *µ* and *σ* values for each bin, *n*. The trained model can be applied to a set of user defined regions, 1… *K*, to evaluate relative mutation burden. This testing is associated with a set of p-values, p, for each of the K regions. The p-values from multiple regions may be combined using Fisher’s method.

## Results

We designed our NIMBus model based on statistical and biological problems inherent in our data, and then we demonstrated our model’s many applications. Compared to the Poisson distribution, the negative binomial model better accounts for the heterogeneity and overdispersion in the mutation rate across diseases and samples. Local mutation rates are also affected by genomic features such as endogenous DNA damage and chromatin organization. NIMBus addresses both of these concerns by utilizing a negative binomial regression and also by adjusting the local background mutation rate based on the genomic context. The covariates used in our model achieved the best prediction accuracy when the tissue type was matched, however when matching was not possible, pooling the tissue types increased power. Because of the availability of established knowledge surrounding coding regions, we first used NIMBus to identify significantly mutated coding regions. Then, we also tested our model on KEGG pathways aimed at pinpointing significant pathways comprised of these coding regions. Additionally, we applied NIMBus to noncoding regions and identified both well documented and novel burdened genes. Lastly, we benchmarked our result against other methods [23, 24] and compared it to a list of empirically supported cancer genes from COSMIC [25].

### Heterogeneity from various sources leads to large overdispersion in mutation counts data

First, it is known that there is significant mutation rate heterogeneity across diseases and samples. For this reason, it is usually incorrect to assume a homogeneous mutation rate per nucleotide or to consequently use binomial tests to calculate P values. To demonstrate this, we collected WGS variants from 649 cancer patients and 7 cancer types (Fig. S1). In our data, the median number of variants was as low as 70 in Pilocytic Astrocytoma (PA) and as high as 21,287 in Lung adenocarcinoma (LUAD). Even within the same cancer type, mutation counts vary dramatically from sample to sample (lowest at 1,743 and highest at 145,500 in LUAD, Fig. 2A). In addition, there are also large regional mutation rate differences within the same sample (Fig. S2). Therefore, distributions based on constant mutation rate assumption usually fit poorly to mutation count data (Fig. 2B, dashed lines with +, Fig. S3). In light of these issues, we utilized a two-parameter negative binomial distribution to further capture the over-dispersed nature of mutation counts data, which improves fitting to real data significantly (dashed lines with star in Fig. 2B).

**Figure 2.**
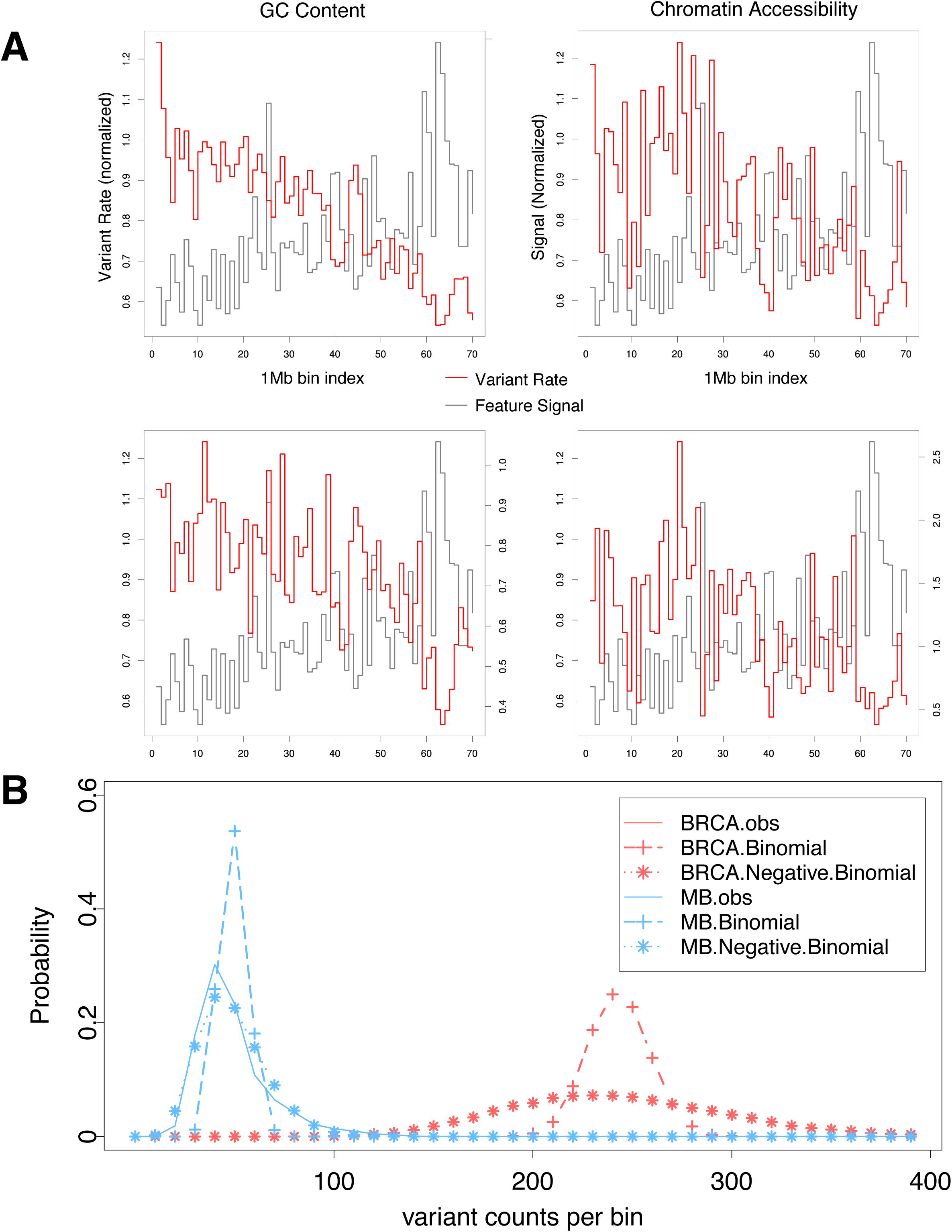
(A) Disease and sample mutation rate heterogeneity; (B) improved fitting by negative binomial distribution of mutation counts in 1mb bins in breast cancer (BRCA) and Medulloblastoma (MB).

### Local mutation rate is confounded by many genomic features

It has been reported that local mutation rates are associated with many well-known genomic features, such as mRNA expression, GC content, replication timing, and chromatin organization [11]. We found that the WGS data in our datasets also demonstrated similar characteristics. For example, Fig. S4 shows how mutation counts at a 1mb resolution (the first 70 bins on chromosome 1) are correlated with several genomic features.

Somatic mutation rate has been reported to be confounded by several genomic features [7-9]. We examined two such genomic features: endogenous DNA damage and chromatin organization. Endogenous DNA damage, such as oxidation and deamination, can affect single-stranded DNA during replication. The accumulative damage effect in the later replicated regions will result in increased mutation rate. We have observed a similar trend in our data. For example, the Pearson correlation between normalized mutation counts and replication timing values in BRCA is as high as 0.67 in the first 70 1mb bins (Fig. S4A).

Another example is that the chromatin organization, which arranges the genome into heterochromatin- and euchromatin-like domains, has a dominant influence on regional mutation rate variation in human somatic cells [11]. Consistently, we also find that mutation counts are significantly associated with the DNase-seq signal (Pearson correlation=-0.61, P=1.52x 10^−8^, Fig S4B). Therefore, it is important to accurately estimate local background mutation rate for mutation burden analysis.

### Negative binomial regression precisely estimates local mutation rates by correcting the influence of many genomic features

#### (A) Features in matched tissues usually provide best prediction accuracy but features in unmatched tissue still help

It has been reported that the most accurate local mutation rate prediction can be achieved by using features from matched tissue [9]. Hence, we specifically selected variants in two distinct cancer types, breast cancer (BRCA) and medulloblastoma (MB), and predicted their local mutation rates with a few features from matched (or loosely matched) and unmatched tissues (Table S2). Relative error, defined as the normalized difference of observed and predicted value (Equation 1), was used to assess model performance. Consistent with previous analyses, we find that features in matched tissues usually outperform those from unmatched tissues. For example, the relative error is only 0.128 by using breast tissue related features to predict BRCA mutation rates, which is noticeably smaller than an error of 0.195 when using brain related features (Table S3). Similarly, brain related features have more predictive power compared to breast related ones for MB mutation rates (error of 0.135 VS. 0.183).

Specifically, we represented mutations rates in BRCA and MB as 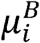 and 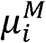 for the *i*^*th*^ bin 1mb bin. 7 genomic features in breast related features were extracted from REMC, including DNASeq, H3K27me3, H3K36me3, H3K4me3, H3K9me3, mRNA-seq and methylation data (features starting with B_ in Fig. S6A), denoted by 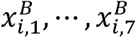. Similarly, we also used 8 features in brain related tissues for MB denoted by 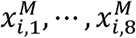 (H3K27me3, H3K27ac, H3K36me3, H3K4me1, H3K4me3, H3K9me3, mRNA-seq and methylation, features starting with M_ in Fig. S6A). We found that these features were correlated both within and across tissues (as shown in the correlation plot in Fig. S6A).

To compare the performance of regressions using (loosely) matched and unmatched tissues, four regression models can be run as shown in Table S2. The scatter plots of the observed and predicted values were given in Fig. S6B. To compare model performance, we defined the relative error 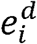 as

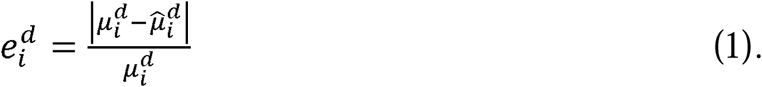

Relative errors for these four models were given in Table S3.

However, biologically meaningful tissue matching remains challenging and usually is not an optimal process for researchers without enough domain knowledge. Specifically, if samples of distinct hidden subtypes were pooled together for a certain disease, tissue matching would be more difficult. Furthermore, even after the optimally matched tissue has been identified, we frequently need to handle missing features in that tissue. We noticed that many genomic features are correlated both within and across tissues (correlation plot in Fig. S6A), which leads to suboptimal but still decent regression performance (scatter plots given in Fig. S6B). This is extremely helpful when processing WGS from diseases without matched features. For example, there are no prostate related features in REMC, but features in other tissues still help to estimate the local mutation rates.

#### (B) Pooling features from multiple tissues significantly improves local background mutation rate prediction

In light of the correlated nature of covariates, especially those epigenetic features [9, 26], we first performed principal component analysis (PCA) on the covariate matrix to address the multicollinearity problem during regression. The correlation of each principal component (PC) with the mutation counts data varies significantly across different cancer types (boxplots in Fig. S7B). For example, the first PC demonstrates a Pearson correlation of 0.653 in LICA, which is much higher than 0.352 in PRAD. Therefore, it is necessary to run a separate regression model for each cancer type.

Since numerous PCs have been shown to be associated with mutation rates, we tried to investigate the joint effect of multiple PCs to predict the local mutation rates. Particularly, for each cancer type, we first ranked the individual PCs by their correlations with mutation rates, and then selected the top 1, 30, and 381 PCs to estimate the local mutation rate. Fig. 3A shows that using more PCs can noticeably boost prediction accuracy in all cancer types. For example, in BRCA the Pearson correlation is only 0.472 if 1 PC is used in regression, but rises to 0.655 and 0.709 if 15 and 30 PCs are used respectively. The correlation increases to 0.818 after using all 381 PCs. As a result, in all of the following analyses, we used all 381 PCs for accurate local mutation rate estimation.

**Figure 3.**
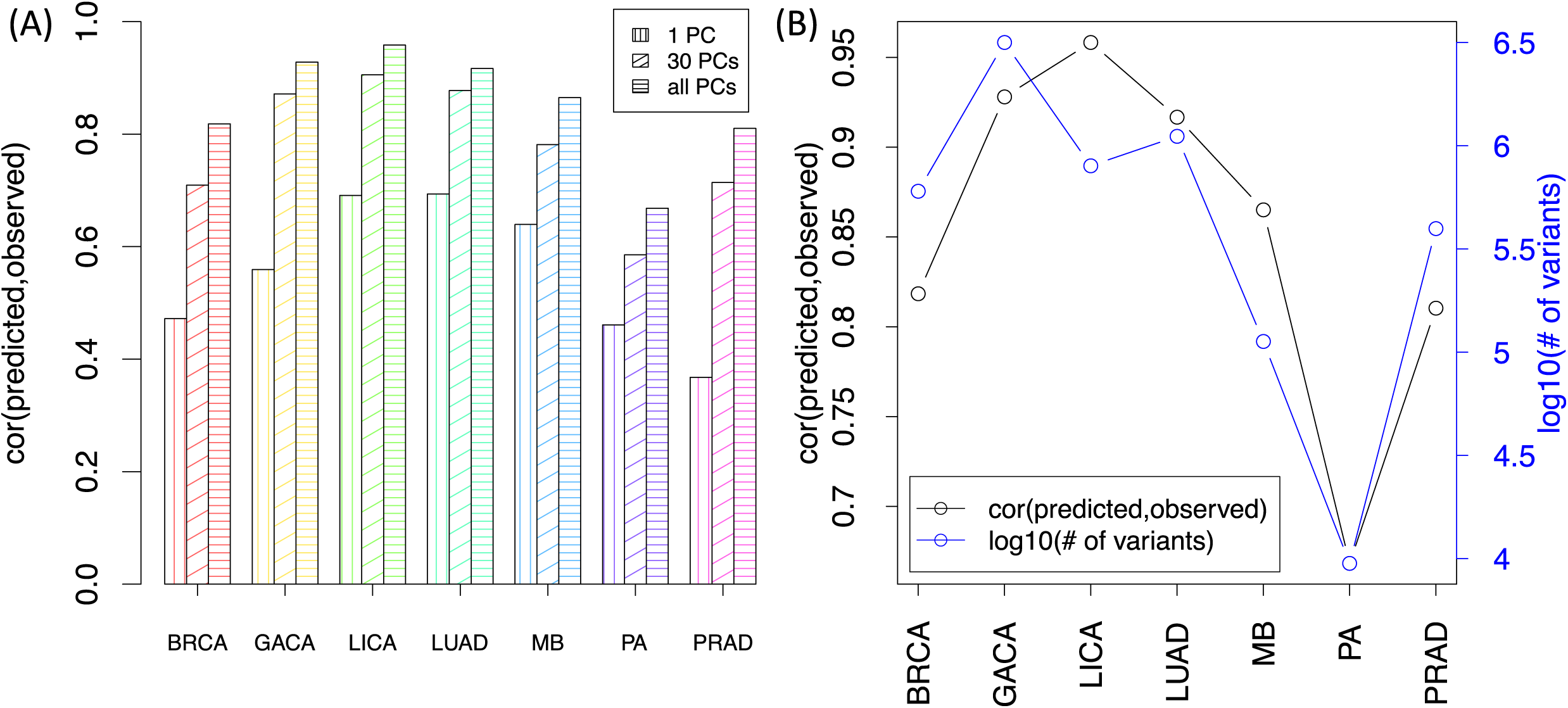
(A) Regression performance by correcting different number of PCs; (B) Regression performance vs. total number of variants used in all cancer types

As shown in Fig. 3B, we achieved good prediction accuracy through regression against all PCs of the covariate matrix in all cancer types. The Pearson correlations of the observed mutation counts and the predicted 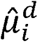 vary from 0.668 in PA to 0.958 in LICA. Scatter plots are given in Fig. S8.

It has been reported that many genomic signal tracks demonstrate noticeable correlations across features and tissues [26]. Hence, we first centered and scaled the covariate matrix *X* and then performed PCA on it to obtain *X*′. The cumulative proportion of variance explained by the PCs was given in Fig. S7 A. As expected, there is lots of redundancy in the covariate table. The first PC may explain as much as 55.69% of variance, and it takes up to 15 and 106 PCs to capture 90% and 99% of variance.

We also calculated the Pearson correlation of PC *j* with mutation counts in cancer type *d* as _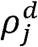_. Then the absolute correlation value _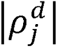_ was averaged over different cancer types as _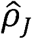_ to rank the PCs. The top 20 PCs with highest _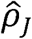_ were selected and boxplot for each of the PCs was given in Fig. S7 B.

For each cancer type, we tried to predict the local mutation rate by correcting the covariate matrix after PCA projection. The Pearson correlation of the predicted and observed mutation rates are given in Fig. S8. It is worth mentioning that although there are no features matching prostate tissue in REMC, we can still achieve a very high correlation of 0.81 with the help of 381 unmatched but correlated features. This indicates that our model can still provide acceptable performance even when somatic WGS of a disease is given without optimally matched covariates.

In addition, the number of available variants obviously affects prediction performance, though it is not the only factor. As shown in Fig. 3B, the limited number of variants, such as those in the quiet somatic genomes of pilocytic astrocytoma (PA), can usually restrict our prediction precision (lowest correlation at 0.668 among 7 cancer types). However, other factors, such as the number of effective covariates, quality of mutation calls, and molecular similarity of pooled samples of the same disease can also influence the prediction performance considerably. For instance, although there are fewer variants in MB than those in BRCA, our regression for MB still outperforms that of BRCA (0.865 vs 0.818, Fig. 3B).

### Coding region calibration for NIMBus

#### (A) Single gene target region analysis

Since coding regions have been investigated in more detail than the noncoding regions, we first applied NIMBus on coding regions. First, we extracted coding regions from the GENCODE annotation v19 and ran NIMBus on both real and simulated datasets (details in Methods “Coding region annotation” and “Simulated variants for all cancer types” sections). We found that in all cancer types analyzed, NIMBus effectively controlled P value inflation compared to the method mentioned in [4]. For example, in LUAD the P values for real data fall on the diagonal with the uniform P values, apart from a few outliers that represent the true significant signals (black dots on the right side in Fig. 4). After P value adjustment using the Benjamin–Hochberg method, only 11 genes are reported as mutated in LUAD, while none were discovered on the simulated data (orange dots in Fig. 4). On the other hand, the method using a constant mutation rate assumption reported 6,023 genes to be significantly mutated, indicating severe P value inflation.

**Figure 4.**
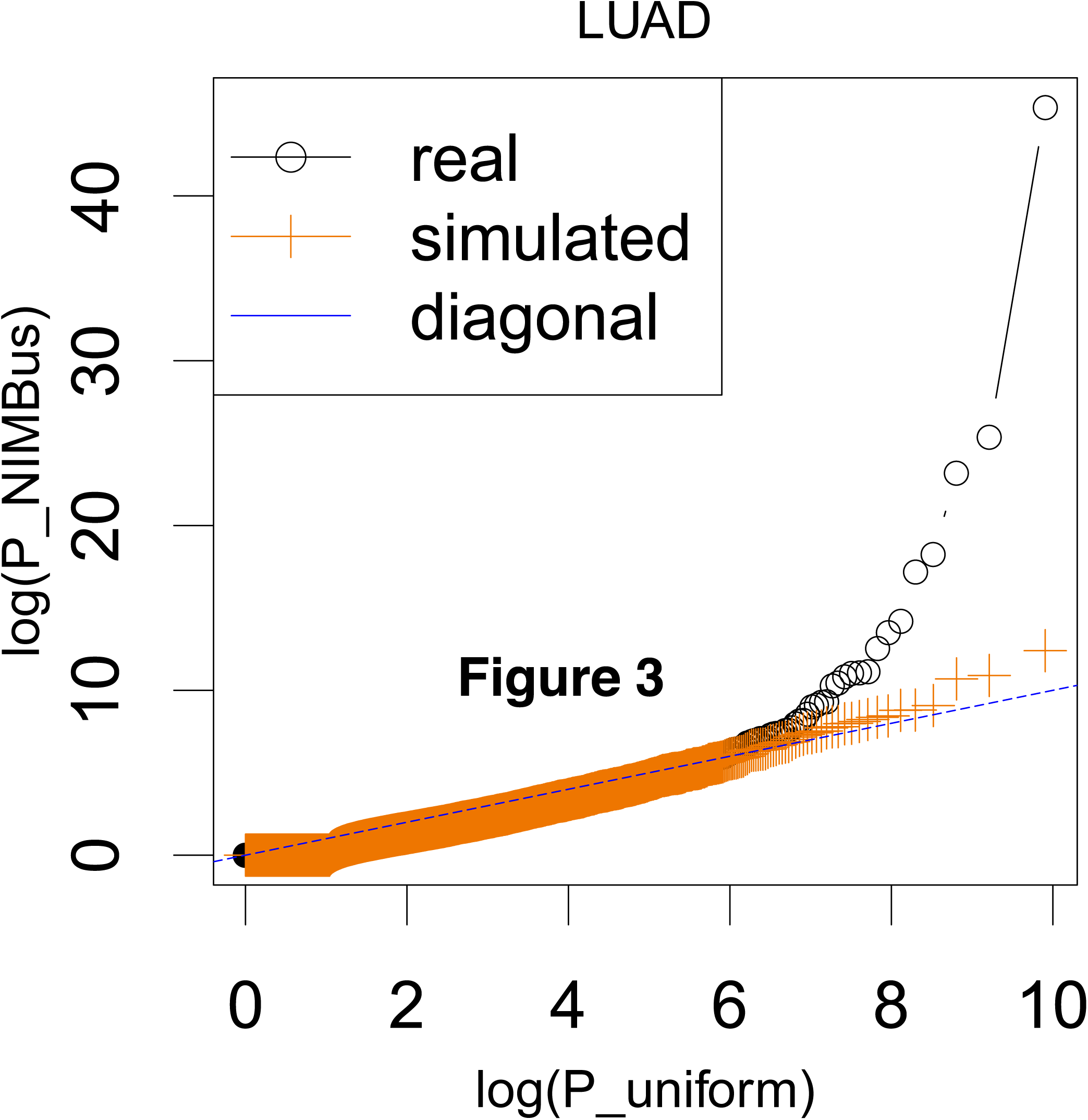
Q-Q plots of P values of real and simulated WGS data.

We also used Fisher’s method to combine P values from all cancer types. In total, 15 genes were discovered to be significantly mutated. Twelve of them are well documented as related with cancer progression. The top genes are shown in Table 1 and their PubMed ID is given in the last column for reference. These results showed that NIMBus is able to find sensible mutational hotspots as cancer drivers.

**Table 1.**
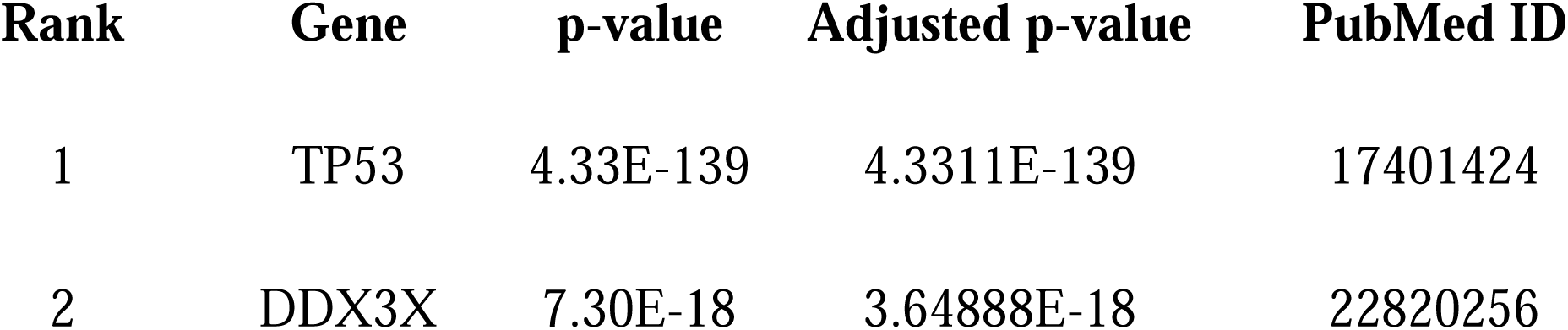
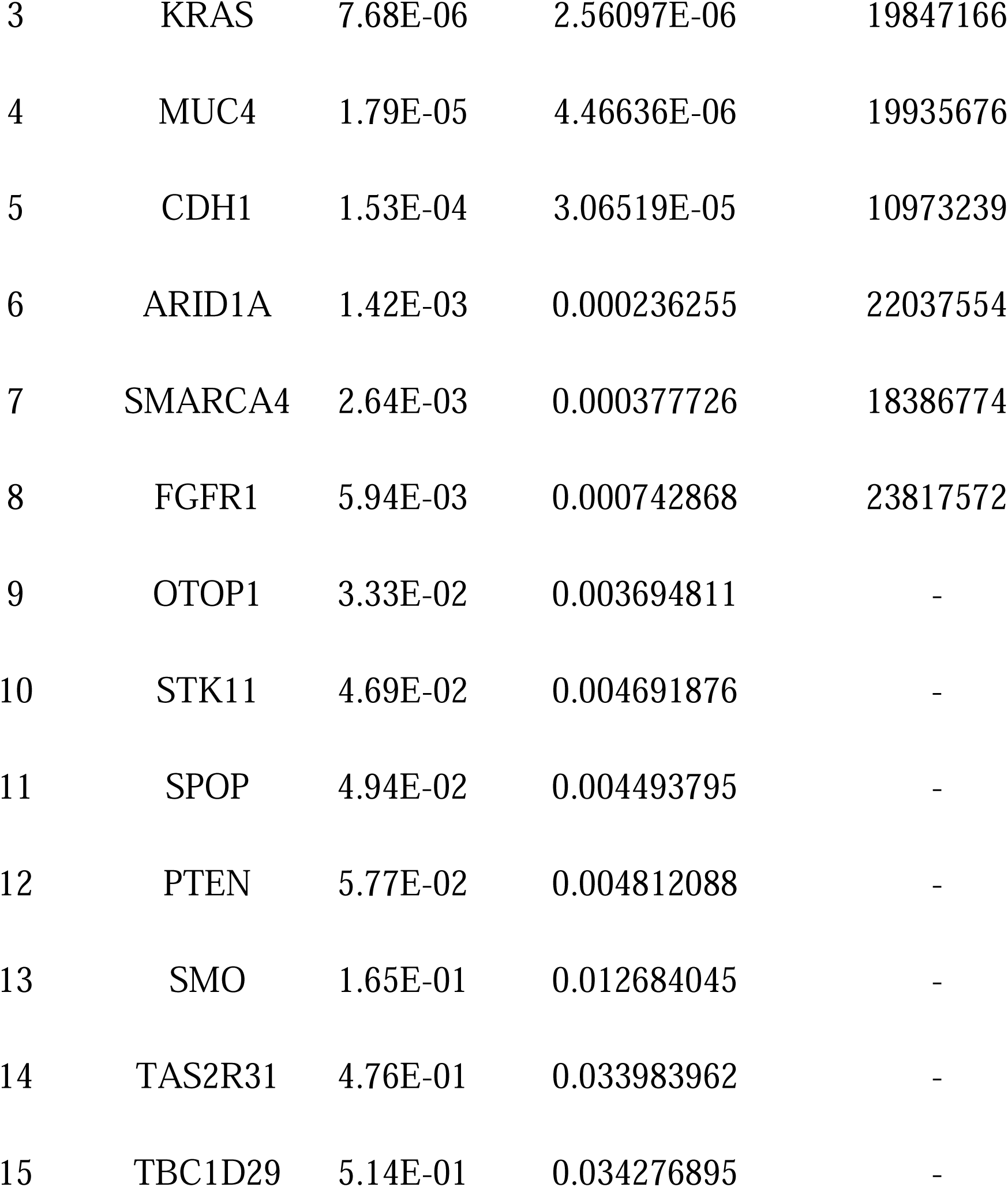
Top genes after p-value combination

| **Rank** | **Gene** | **p-value** | **Adjusted p-value** | **PubMed ID** |
| --- | --- | --- | --- | --- |
| 1 | TP53 | 4.33E-139 | 4.3311E-139 | 17401424 |
| 2 | DDX3X | 7.30E-18 | 3.64888E-18 | 22820256 |
| 3 | KRAS | 7.68E-06 | 2.56097E-06 | 19847166 |
| 4 | MUC4 | 1.79E-05 | 4.46636E-06 | 19935676 |
| 5 | CDH1 | 1.53E-04 | 3.06519E-05 | 10973239 |
| 6 | ARID1A | 1.42E-03 | 0.000236255 | 22037554 |
| 7 | SMARCA4 | 2.64E-03 | 0.000377726 | 18386774 |
| 8 | FGFR1 | 5.94E-03 | 0.000742868 | 23817572 |
| 9 | OTOP1 | 3.33E-02 | 0.003694811 | - |
| 10 | STK11 | 4.69E-02 | 0.004691876 | - |
| 11 | SPOP | 4.94E-02 | 0.004493795 | - |
| 12 | PTEN | 5.77E-02 | 0.004812088 | - |
| 13 | SMO | 1.65E-01 | 0.012684045 | - |
| 14 | TAS2R31 | 4.76E-01 | 0.033983962 | - |
| 15 | TBC1D29 | 5.14E-01 | 0.034276895 | - |

#### (B) Mutation burden of KEGG pathways

Using the KEGG pathway dataset, consisting of 288 unique pathways, we performed a network mutation burden test on each pathway for each cancer type to discover significantly mutated pathways. We found that of the seven cancer types analyzed, four cancer types exhibited significantly mutated KEGG pathways (*p*_*adj*_ < 0.05). In particular, we found 5 significant pathways in BRCA, 5 in LICA, 10 in GACA, and 3 in LUAD. No significant pathways were found in MB, PA, or PRAD. The significant pathways and their associated cancer types are seen in Table 2, as well as the Benjamin-Hochberg adjusted *p*-value. The significant pathway list includes pathways associated with the p53 signaling pathway, apoptosis, and cell growth – which are known to be associated with cancer. In addition to these well-studied pathways, we were able to discover many novel pathways, including other signaling and disease-associated pathways. These results demonstrate a novel way to use NIMBus as a way to conduct mutation burden tests in biologically meaningful networks in the genome.

**Table 2.**
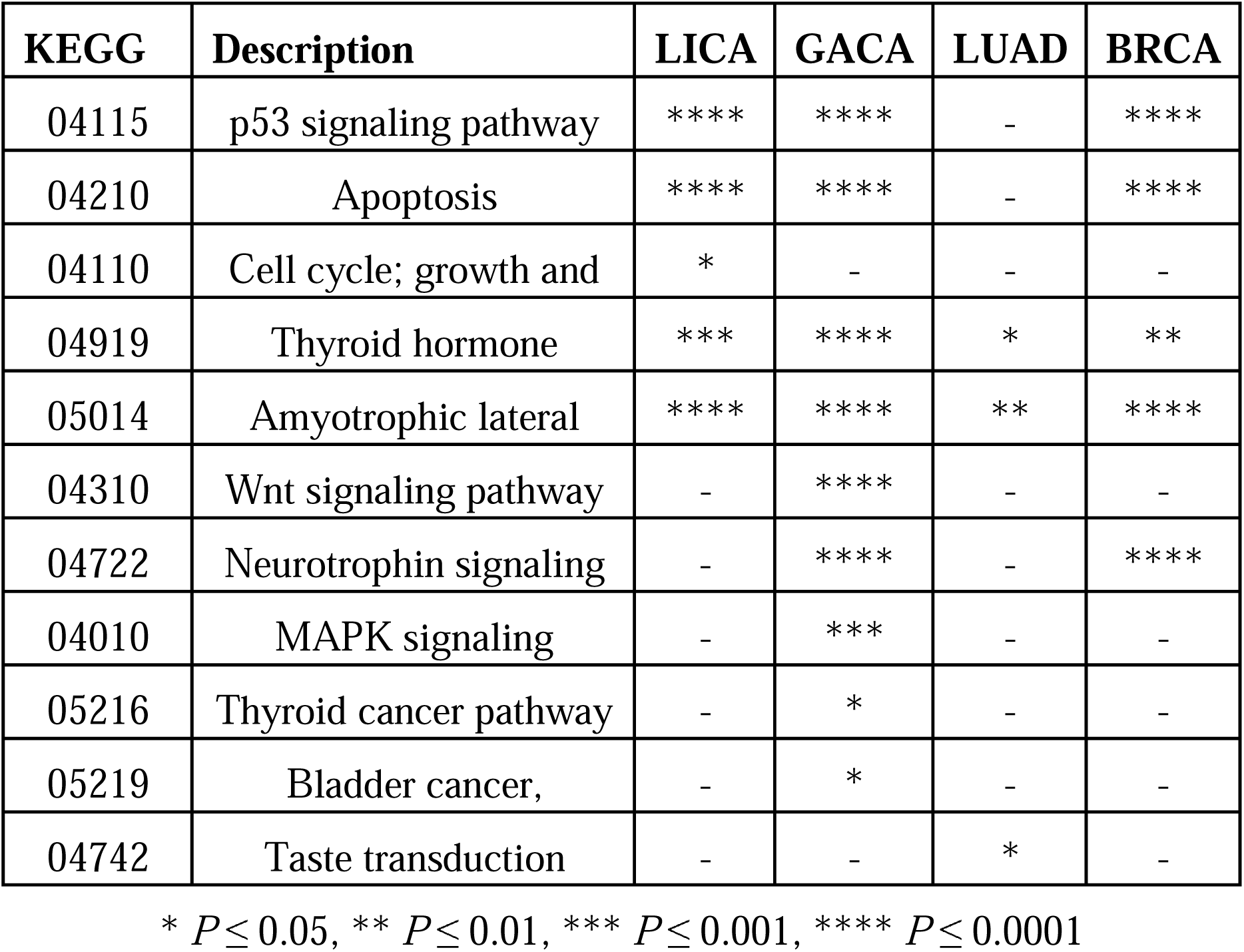
Significant pathways and *P* values

| KEGG | Description | LICA | GACA | LUAD | BRCA |
| --- | --- | --- | --- | --- | --- |
| 04115 | p53 signaling pathway | **** | **** | - | **** |
| 04210 | Apoptosis | **** | **** | - | **** |
| 04110 | Cell cycle; growth and | * | - | - | - |
| 04919 | Thyroid hormone | *** | **** | * | ** |
| 05014 | Amyotrophic lateral | **** | **** | ** | **** |
| 04310 | Wnt signaling pathway | - | **** | - | - |
| 04722 | Neurotrophin signaling | - | **** | - | **** |
| 04010 | MAPK signaling | - | *** | - | - |
| 05216 | Thyroid cancer pathway | - | * | - | - |
| 05219 | Bladder cancer, | - | * | - | - |
| 04742 | Taste transduction | - | - | * | - |

\* $P \leq 0.05$ , \*\* $P \leq 0.01$ , \*\*\* $P \leq 0.001$ , \*\*\*\* $P \leq 0.0001$

### NIMBus discovered a list of mutated noncoding regions from cancer WGS data

As a fair comparison to our NIMBus model, the global and local models were used on the same data to identify mutational hotspots. First, the global model assumes the per nucleotide mutation rate is constant across the genome and different individuals. Hence the mutation counts within the test region could be considered as a Binomial distribution. Then, the local model uses a Binomial regression against the same set of covariates to compensate regional mutational heterogeneity, but ignores the heterogeneity across individuals. On the other hand, our NIMBus model captures mutational heterogeneity arising from both different individuals and regions from the genome, which allows more flexibility of mutation counts modeling (details see the method section).

We applied NIMBus on WGS variant calls for all seven cancer types to predict the individual somatic burden P values, and compared these results to those from global and local Binomial models (details in the Methods “WGS variants data used” section). Similar to the results in the coding region analysis, both global and local Binomial models generated too many burdened regions in all noncoding annotation categories, as evidenced by the poor fitting in Fig. 2B. For example, in liver cancer after P value correction, NIMBus identified 21 promoters as highly mutated, while local and global binomial models identified 79 and 641, respectively. Hence, our negative binomial assumption in NIMBus effectively captured the overdispersion and controlled the number of false positives. To further demonstrate this, we provided the Q-Q plots of P values in promoter regions for all seven cancer types in Fig. 5B as a quality check. In theory, if no significantly burdened regions are detected, the P values should follow uniform distribution. As shown in Fig. 5B, the majority of our P values follow the uniform assumption, with the exception of a few outliers representing true signals, indicating reasonable P value distributions for all cancer types. BRCA and GACA, as well as PRAD and LUAD, have a greater number of outliers than other cancer types. This may relate partially to statistical power – these cancer types have a greater number of mutations and more patients than those cancer types without the same number of outliers. In addition to the QQ plots in Fig. 5B, we also looked at the proportion of significantly burdened promoter regions for each cancer type. We found that GACA has more than twice the number of burdened promoter regions as other cancer types. In addition to statistical power to measure this burdening, this may also relate to underlying differences in tumor biology.

**Figure 5.**
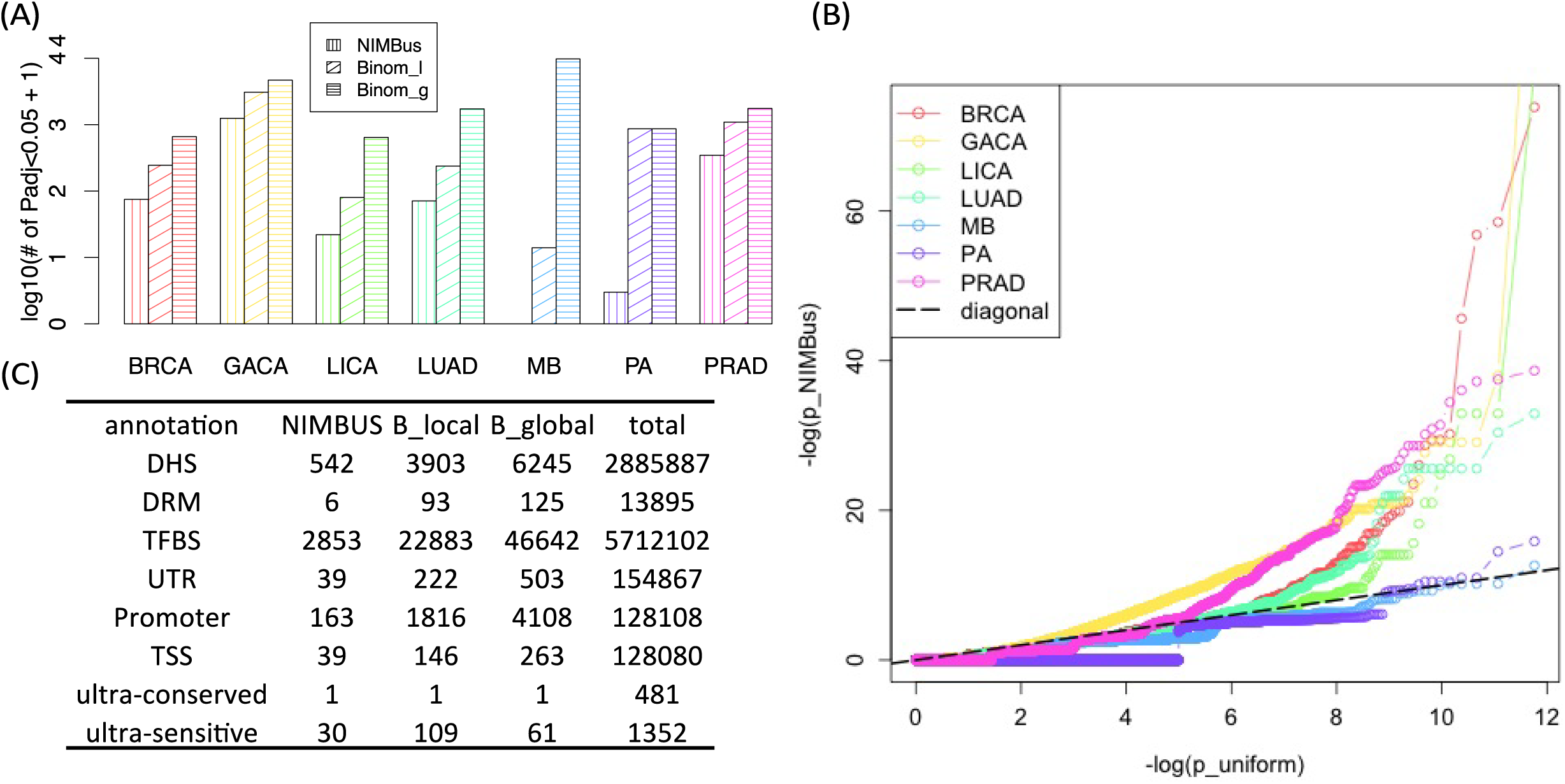
(A) Number of overly mutated promoter regions in all cancer types; (B) Q-Q plots of P values for promoter regions; (c) total number of burdened regions in all noncoding annotations after merging P values from 7 cancer types. B_local: local Binomial Model, B_global: global Binomial Model, DRM: Distal Regulatory Module, DHS: DNase hypersensitivity site, TFBS: Transcription factor binding site, UTR: Untranslated region, Promoter: 2500 nucleotides (nt) upstream of the 1^st^ nucleotide of GENCODE transcripts, TSS: Transcription Start Site. 100 nucleotides upstream of the 1^st^ nucleotide of GENCODE transcripts, Ultraconserved region: region under positive cross-species selection in mammals, Ultrasensitive region: region with a greater than expected fraction of rare variants.

To summarize the mutation burdens from all cancer types, we used Fisher’s method to calculate the final P values for all three models. Similar to P values from a single cancer type, the combined P values are severely inflated in both global and local Binomial models, but are rigorously controlled by NIMBus (table C in Fig. 5). For example, NIMBus reported only 39 transcription start sites (TSS) as burdened, compared to 164 and 263 for the other two methods.

Additionally, out of the 39 TSS elements, several of them have been experimentally validated or computationally predicted in other work to be associated with cancer. For instance, TP53 is a well-studied tumor suppressor gene that is involved in many cancer types, and the combined P value for the TP53 TSS is ranked second in our analysis (P=4.26× 10^−14^) [27]. Another TSS element found to be significantly burdened is LMO3, which interacts with the tumor suppressor TP53 and regulates its function. LM03 ranked fourth in our analysis (P=3.25× 10^−13^) [28]. Similar to previous reports, we also identified the AGAP5 TSS site as a mutation hotspot, ranking third (P=7.07× 10^−14^) in our analysis [28]. Another important example is the TSS site in TERT, which ranked fifth in our results (P=1.55× 10 ^−14^) and has been experimentally validated to be associated with multiple types of cancer progression [19-21]. The discovery of such results shows that NIMBus may contribute to mutation driver event discovery in genetic diseases.

To further extend our analysis of non-coding mutational burdening, we examined transcription factor binding sites within promoter regions with evidence of functional activity via co-location with DNase I hypersensitive sites (DHSs). This analysis was performed on a per-cancer type basis, with assays matched to cancer types (Table S5). In total, 14 cancer types were examined. For 7 of the cancer-types with available RNA-seq expression in ENCODE, gene expression was used to limit the analysis to TFs with non-negligible expression (>1 FPKM). In total, after multiple-testing correction, 1450 noncoding regions were identified with significant mutation burdening. Although significantly burdened sites were identified for all cancer types analyzed, certain cancer types contained disproportionate numbers of burdened promoter/TF sites. Skin-Melanoma (401 burdened sites), Myeloid-MPN (296 burdened sites), and Lymph-BNHL (274 sites), were the cancer types with the greatest number of non-coding promoter/TF regions identified. The promoter/TFBS regions of common cancer-associated genes were identified as overburdened, such as TERT in Liver-HCC and TP53 in Lung-SCC.

We compared the burdened noncoding regions identified by NIMBus to those identified using another method, OncodriveFML [24]. Across 14 cancers, OncodriveFML identified a total of 249 unique burdened noncoding regions, while NIMBus identified a total of 1,380 unique burdened noncoding regions. Twenty-three genes were shared between both methods, including STAT3 and MYC. Notably, OncodriveFML did not identify the noncoding regions of TERT or TP53 to be significantly burdened.

We also compared the regions identified through this method, to an independently derived set of non-coding driver mutations identified by the ICGC/TCGA Pan-Cancer Analysis of Whole Genomes Consortium [23]. The PCAWG Consortium tested six different types of noncoding regions (3’UTR, 5’UTR, enhancers, promoters, long non-coding RNA genes (lncRNA), and promoters of lncRNA genes). Overall, NIMBus found the noncoding regions of 1,380 unique genes to be burdened across 14 cancers, while the PCAWG Consortium’s noncoding driver analysis found a total of 29 genes to be burdened across 12 cancers. Eighteen burdened genes were shared in both analyses, but NIMBus identified 1,362 additional unique genes that the PCAWG Consortium did not. The shared genes included the well-studied TERT gene. While the PCAWG Consortium found TP53 to be mutated in its coding region, their noncoding analysis did not find it to be mutationally burdened. Using NIMBus we find TP53 to be significantly burdened in both its coding and noncoding regions, which is supported heavily in literature [29].

We used the Cancer Gene Census as compiled by COSMIC (Catalogue of Somatic Mutations in Cancer) to further analyze NIMBus’s performance [25]. Cancer genes were either classified as Tier 1 (possessing strong evidence) or Tier 2 (possessing developing evidence). Of the 18 burdened genes that were identified by NIMBus and the PCAWG Consortium, one gene (TERT) was identified as a Tier 1 gene and one gene (MALAT1), a lncRNA gene, was identified as a Tier 2 gene [30]. While NIMBus identified a total of 56 Tier 1 genes and 13 Tier 2 genes, the PCAWG Consortium identified 2 Tier 1 genes and 1 Tier 2 gene; and OncodriveFML identified 37 Tier 1 genes and 1 Tier 2 gene. Among the Tier 1 genes identified by NIMBus, CXCR4 is a chemokine receptor that interacts with the chemokine molecule CXCL12 [31]. The CXCR4/CXCL12 pathway has substantial literature support that establishes the role of the CXCR4 gene in cancer [32, 33]. Another Tier 1 gene identified by NIMBus, ERBB2, is a transmembrane tyrosine kinase receptor that is often overexpressed in cancer [34]. Anti-ERBB2 antibodies (under the generic name trastuzmab) can be used to treat breast and gastric cancer [35, 36]. Our benchmarking results are summarized in an Excel spreadsheet on our NIMBus Github website.

By comparing our results to those generated from well-known, previously existing methods, we find that NIMBus not only robustly pinpoints already well documented findings, but also identifies additional burdened genes with both developing and existing experimental support.

## Discussion

Thousands of somatic genomes are now available due to the fast development of whole genome sequencing technologies, providing us with increasing statistical power to scrutinize the cancer mutation landscape. At the same time, thanks to the collaborative efforts of large consortia, such as REMC and ENCODE, tens of thousands of functional characteristic experimental results on human genomes have been released for immediate use to the whole community. Hence, integrative frameworks are of urgent need in order to explore the interplay between WGS data and these functional characteristic data. It will not only be important to accurately search for mutational hotspots as driver candidates for complex diseases but also to better interpret the underlying biological mechanisms of diseases for clinicians and biologists.

In this paper, we proposed a new integrative framework called NIMBus to analyze cancer genomes. Due to the heterogeneous nature of various somatic genomes, our method treated the individual mutation rate as a gamma distributed random variable to mimic the varying mutational baseline for different patients. Resultantly, it modeled the mutation counts data using a two-parameter negative binomial distribution, which improved data fitting dramatically as compared to previous work (Fig. 2B). It then uses a negative binomial regression to capture the effect of a widespread list of genomic features on mutation processes for accurate somatic burden analysis.

Unlike previous efforts, which use very limited covariates to estimate local mutation rate in very qualitative way, we explored the whole REMC and ENCODE data and extracted 381 features that best describe chromatin organization, expression profiling, replication status, and context effect in all possible tissues to jointly predict the local mutation rate at high precision. In terms of covariate correction, NIMBus demonstrated three obvious advantages: 1) It incorporates the most comprehensive list of covariates in multiple tissues to achieve accurate background mutation rate estimation; 2) It provides an integrative framework that can be extended to any number of covariates and successfully avoids the high dimensionality problem of other methods [5, 7], which is extremely important due to the rapidly growing amount of available functional characteristic data available and the drop in cost of sequencing technologies; 3) It automatically utilizes the genomic regions with the highest credibility for training purposes, so users do not have to be concerned about performing carefully calibrated training data selection and complex covariate matching processes.

The length of training bins *l* is an important parameter for NIMBus. On one side, a shorter bin size will be advantageous in the P value evaluation as it can remove the mutational heterogeneity across regions more effectively at a higher resolution. On the other side, a smaller *l* sometimes will result in worse mutation rate prediction performance for two reasons. First, sensible mutation rate quantification is necessary in each single bin for the regression purpose. However, somatic mutations are usually sparsely scattered across the genome due to limited number of disease genomes available at the moment. In the extreme case, when *l* is so small that most bins have zero mutations, it is difficult for the regression model to capture the relationship between mutations and covariates. Second, some of the covariates are only reported to be functional on a large scale [11], so reducing *l* will not necessarily boost prediction precision. Optimal bin size selection is still a challenging problem that needs further case-by-case investigation. In our analysis, we used a 1 Mb bin size for all cancer types.

Noncoding regions represent more than 98% of the whole human genome, and are investigated less mainly due to limited knowledge of their biological functions. NIMBus is also designed to explore the most comprehensive collection of noncoding annotations. Therefore, it collects the up-to-date, full catalog of noncoding annotations of all possible tissues from ENCODE and our previous efforts from the 1000 Genomes Project. Additionally, it further customizes these annotations specifically for somatic burden analysis. All these integrated internal annotations of NIMBus can be either tested for somatic burden or used to annotate the user specific input regions.

We applied NIMBus to 649 cancer genomes of seven different types collected from public data and collaborators. The burden test P values for each cancer type were deduced and Fisher’s method was used to calculate the combined P values. We first evaluated the performance of NIMBus on coding regions, which have been investigated with much detail by researchers. Many well-documented cancer associated genes were discovered by NIMBus (Table 1 and Table S4). We also repeated the same analysis on a simulated dataset and found no significant genes. These results demonstrate that NIMBus is able to find overly mutated genes effectively while rigorously controlling false positives. It should be mentioned that one limitation of this analysis is the limited statistical power due to a lack of WGS data. This in turn results in appropriately conservative predictions. However, with the increasing availability of more WGS data and advances in sequencing technology, finding a more complete list of heavily burdened genes will become possible.

In addition to single gene burden analysis tests, we were able to detect significantly mutated KEGG pathways, including the TP53 signaling pathway and apoptosis pathway, both of which are implicated in cancer progression. We also examined the burdening of TFBS located in promoter regions that intersect with DHSs. 1450 TFBS/promoter regions were identified with disproportional mutation rates. A subset of these regions was also found to affect downstream gene expression when mutated. This approach identified novel, gene-regulatory regions that may affect cancer development and also provides an example of how the NIMBus method may be extended to examine gene networks. The adaptability of NIMBus to analysis of gene networks may prove useful in determining significantly mutated regions of the genome that are not physically adjacent. Though NIMBus is able to determine some well-known cancer associated pathways that are heavily burdened, there still remains some challenges in interpreting other pathways. One reason may be due to constrained availability of pathway annotations, which may result in false positives. Future work could be done here to build and validate other pathways. Additionally, when using Fisher’s method, it is possible that the p-values of the regions or genes in a pathway that are combined are not entirely independent, which could result in some false positives, since we do not know the exact joint distribution of the p-values. This may be an area of future study.

Furthermore, numerous noncoding elements were also reported as significantly mutated (Table C in Fig. 5). Included were some well-known regions, such as the TP53, LMO, and TERT TSS, proving the effectiveness of NIMBus in identifying disease-associated regions.

To some degree somatic variants can be considered as the limit of extremely rare germline variants because they are almost private variants to particular cells. On the contrary, common germline variants have somewhat different characteristics from rare germline ones as they often have low functional impact and are linked to other variants. As the germline variant becomes more rare, the linkage decreases and the functional impact usually increases up to what we observe for somatic variants. Thus, we would expect the methods here to work well for rare germline variants (e.g. *de novo* ones and those confined to small populations)

Scanning the TFBS within regulatory elements allowed us to gain more power than simply analyzing whole regulatory regions. Some identified burdened genes were shared with the PCAWG Consortium, however we also identified a number of additional genes that are classified by COSMIC as possessing strong experimental evidence.

## Conclusions

In summary, NIMBus is the first method that integrates comprehensive genomic features to analyze the mutation burdens of disease genomes. Such external data does not only help to better estimate the background mutation rate for successful false positive and negative control, but also provides the most extensive noncoding annotations for users to interpret their results. It serves as a powerful computation tool to accurately predict driver events in human genetic diseases and potentially identify biological targets for drug discovery.

## Methods

### WGS variants data used

We collected 649 whole genome variant calls from public resources and collaborators. This data set contains a broad spectrum of 7 different cancer types, including breast cancer (BRCA, 119 samples), gastric cancer (GACA, 100 samples), liver cancer (LICA, 88 samples), Lung adenocarcinoma (LUAD, 46 samples), prostate cancer (PRAD, 95 samples), Medulloblastoma (MB, 100 samples), and Pilocytic Astrocytoma (PA, 101 samples) (Figure S1, Table S1). GACA samples were from Wang *et al* [37] and PRAD samples were obtained from our collaborators. The remaining comes from samples published by Alexandrov *et al* [38].

### Local background mutation rate estimation

#### (A) Feature selection

Numerous studies showed many genomic features severely affect the mutation process, and such covariate effect should be removed for somatic burden analysis [7, 11]. We first collected all the signal track files from major histone modification marks, chromatin status, methylation, and mRNA-seq data from the REMC. Signal files were processed in bigWig format at 20nt resolution. Multiple replicates were averaged if available. Since replication timing has been proved to be associated with mutation rate in several articles [5, 7, 11], we also collected 8 replication timing bigWig files from the ENCODE project. Lastly, as researchers have observed elevated mutation rates in regions wither lower GC content in certain diseases, we also include the GC percentage files in our covariate list and generated its corresponding bigWig files.

#### ((B) Human genome gridding and covariate matrix calculation

Different from the calibrated training data selection mentioned in [17], we divided the whole genome into bins with fixed length *l*, such as 1mb, 100kb, 50kb, etc. Only autosomal chromosomes and chromosome X were included in our analysis to remove the gender imbalance in the mutation data or covariates.

Repetitive regions on the human genome are known to generate artifacts in high throughput sequencing analysis mainly due to their low mappability. We downloaded the mappability consensus excludable table used in the ENCODE project from http://hgdownload.cse.ucsc.edu/goldenpath/hg19/encodeDCC/wgEncodeMapability/wgEncodeDacMapabilityConsensusExcludable.bed.gz. Any fixed length bins that overlap with this table would be removed from the training process. We also downloaded the gap regions of hg19 from the UCSC genome browser, which include gaps from telomeres, short_arms, heterochromatin, contigs, and scaffolds. The fixed length bins that intersect with these gap regions were also removed in our analysis. Together these are known as the blacklist regions.

Then, 381 features are extracted from both REMC and ENCODE, and the average signal in the bins is calculated. All the bigWig files generated in step one were used to calculate the average signal using the bigWigAverageOverBed tool for each fixed length bin we generated above. When calculating the GC percentage, if the sequence information is not available at a certain position (such as the Ns), such position will be excluded in the averaging process. In the end, we summarized all the covariates values in each bin into a covariate table, with columns indicating different features and rows representing different training bins. We let *x*_*i,j*_ denote the average signal strength for the *i*^*th*^ bin and *j*^*th*^ covariate, where *i* = 1,..,*n* and *j* = 1,..,*m*.

#### ((C) Use negative binomial distribution to handle mutation count overdispersion

Suppose there are *d* 1,..,*D* different diseases (or disease types) in the collected WGS data, and *s* = 1,..,*s*_*d*_ unique samples, for example different patients, for each disease (or disease type such as liver cancer or lung cancer) *d*. Let 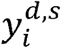 and 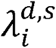 denote the observed mutation count and rate for the *i*^*th*^ bin defined above for sample in disease *d*. In previous efforts, scientists assume that mutation rate 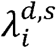 is constant across different regions of the human genome, samples, and diseases, so they have that 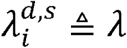 for ∀*i, d, s*. Hence 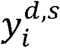 follows a Poisson distribution with the probability mass function (PMF) given in equation (2).

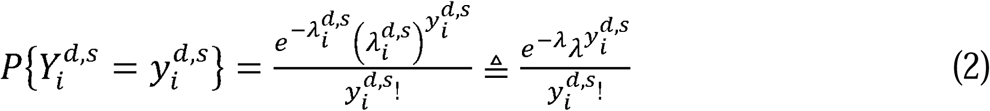

However, somatic genomes are heterogeneous because mutation rates vary considerably among various diseases, samples, and regions of the same genome, severely violating the assumption in equation (2). As a result, fitting of 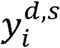 is usually very poor because overdispersion is often observed [5]. Simply assuming a constant mutation rate will generate numerous false positives. Instead, in our model we assume that different 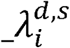 are random variables that follow a Gamma distribution with probability density function (PDF)

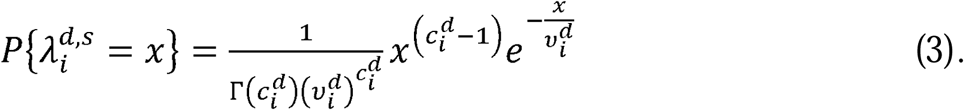

where 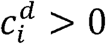 and 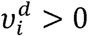. In equation (3), 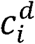 and 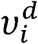 are the shape and scale parameters respectively. Assume that 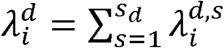 is the overall mutation rate from all samples in bin *i* of disease *d*. Its distribution can be readily obtained through convolution as

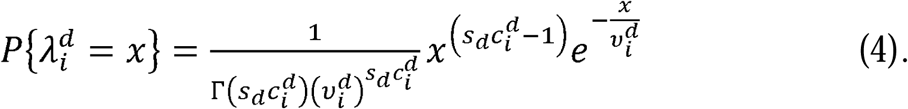

If we let 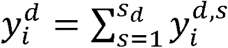 represent the total mutation counts in region *i* from all disease samples, *d*, then the conditional distribution of 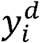 given 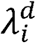 can be written as

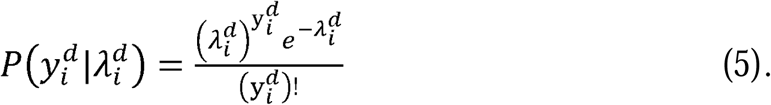

By integrating (4) into (5), the marginal distribution of 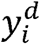 can be denoted as a negative binomial distribution ([39], page 50 in [40]).

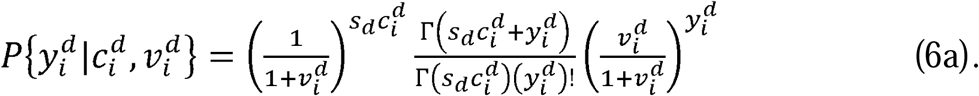

Equation (6a) is the PDF of a negative binomial distribution with 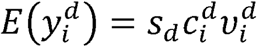 and 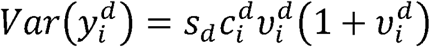. To better interpret (6a), we define 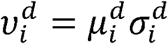 and 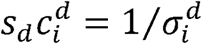. Then equation (6a) can be rewritten as (6b).

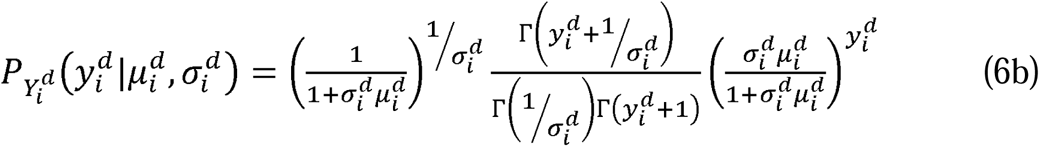

The mean and variance of 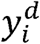 from (6b) can be described as 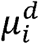 and 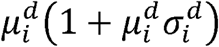 respectively. Our model in equation (6b) is convenient due to its explicit interpretability. First, it assumes that the individual mutation rates are heterogeneous by modeling 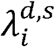 as i.i.d. Gamma distributed random variables. Unlike the constant mutation rate assumption where 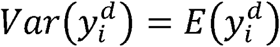, our model captures the extra variance of 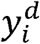 due to population heterogeneity. Our model in (6b) also clearly separates the two main parameters 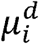 and 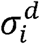 with physically interpretable meanings: the mean and overdispersion, respectively. Here a larger 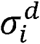 indicates a more severe degree of overdispersion, which is usually due to larger differences in mutation rates.

#### ((D) Accurate local background mutation rate estimation by regression

After modeling 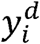 with a negative binomial distribution, we then estimate the local mutation rate by correcting the covariate matrix ***X*** described above. Again *x*_*i,j*_ denotes the average signal strength in the *i*^*th*^ bin and *j*^*th*^ covariate, where *i* = 1,..,*n* and *j* = 1,..,*m*. Because the genomic features in the covariate matrix are correlated and may introduce multicollinearity if directly used in regression, we first apply principal component analysis (PCA) to matrix ***X***. We define ***X*′**to be the covariate matrix after PCA and 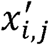 as each element in ***X*′**.

A generalized regression scheme is used here. Suppose *g*_1_ and *g*_2_ are two link functions. We then use linear combinations of covariate matrix ***X*′**′ to predict the transformed mean parameter, 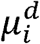, and overdispersion parameter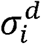, as

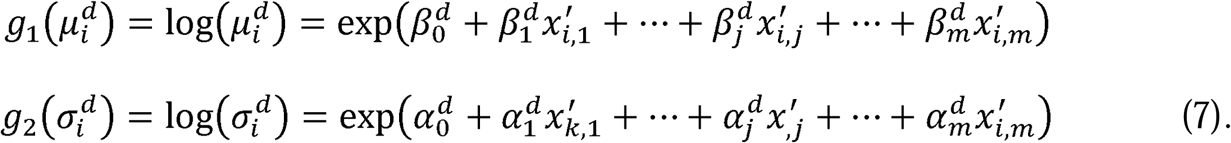

Here we use a log link function for both *g*_1_ and *g*_2_, so the regression model in (7) is a negative binomial regression. Note that x contains 381 genomic features in all available tissues. In the following analysis, we use all features to run the regression in (7) to achieve better performance. The GAMLSS package in R is used to estimate the parameters in (7) as 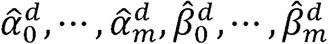. Generally, there are biological reasons to explain how 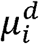 changes with covariates. For example, single-stranded DNA in the later replicated regions usually suffers from accumulative damage resulting in larger 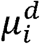. It is more difficult to interpret such a relationship with 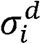. Hence, we simplify equation (7) by assuming 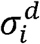 is constant in our real data analysis, meaning the overdispersion parameter *σ*, was modeled as a constant across all bins 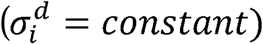for *i* = 1,…,*n*.

After the training process through equation (7) in the main manuscript, the estimates of parameters for negative binomial regression can be represented by 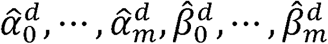. To obtain the optimal local mutation rate for test region *k*, which may be either an internal noncoding annotation such as enhancer or a user-defined element, we should first extend this region into the training bin length *l* centered at the center of test region *k* (blue parts in Fig. S2). Then the covariates values after PCA projection in this extended bin should be calculate as 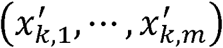. Hence in this scheme, the local mutation parameters should be calculated as

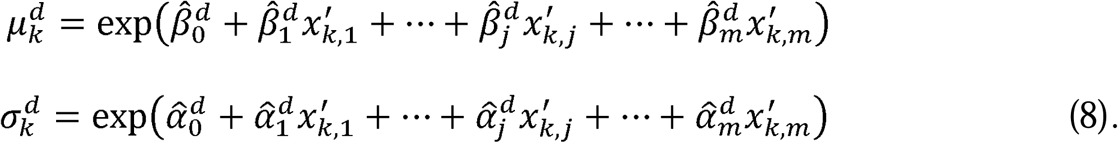

However, in real data analysis there are usually millions of regions to be tested and for each region it needs to process 381 features. Hence, the above optimal scheme is usually\ computational expensive. Here we proposed an approximation scheme to calculate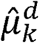 and. 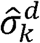. Instead of using covariates for the extended bin centered at target region *k*, we used the values for the nearest training bin 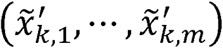 (magenta parts in Fig. S2), and burden tests are performed after length adjustment. Since 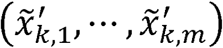has already been pre-calculated during the training process, our approximation scheme significantly reduced the computation burden for tests.

### Somatic burden tests using local background mutation rate

#### (A) Background mutation rate calculation for target regions

Suppose there are *K* regions to be tested. We use the local mutation rate to evaluate the mutation burden. For the *K*^*th*^ target region (*k* = 1,..,*K*), one way of calculating the covariates is to extend it into length *l* (illustrative figure given in Fig. S2). Then we calculate the average signal for feature *j* as *x*_*k,j*_, *j* = 1,..*m* for this extended bin, and after PCA projection let 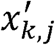 represent the value for the *j*^*th*^ PC. The local mutation parameters 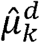 and 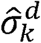 in the extended bin for the *k*^*th*^ target region can be calculated as:

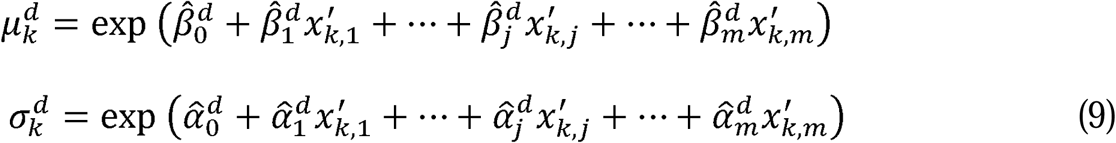

In real data analysis, the length of the *k*^*th*^ test region *l*_*k*_ is much shorter than the length of the training bins (up to 1Mb). Hence 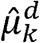 needs to be adjusted by a factor of *l*_*k*_*/l* Then 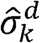 and the adjusted 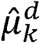 can be used to calculate the disease specific P value, 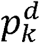. This above scheme is usually computationally expensive because there are usually millions of target regions to be tested. Therefore, we also propose an approximation method to replace the optimal 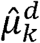 and 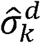 in our analysis (details under Supplemental Fig. S2).

#### (B) Combining P values for multiple disease types

Sometimes it is necessary to analyze several related diseases (or disease types) to provide a combined P value. One typical example is in pan-cancer analysis. In the above section, we calculated the P value for disease/disease type *d* as 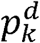 for test region *k* Fisher’s method can be used to combine these P values. Specifically, the test statistic is

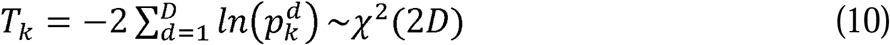

Here *T*_*k*_ follows a centered chi-square distribution with 2*D* degrees of freedom, where *D* is the total number of diseases/disease types. The final P value, *p*_*k*._, can be calculated from *T*_*k*_ To better illustrate how NIMBus works, Fig. 1 gives its workflow.

As a fair comparison to our NIMBus model, the global and local Poisson models were used on the same data to identify mutational hotspots. The global Poisson model assumes the observed mutation counts follow a Poisson distribution and the Poisson rate is constant across the genome and across individuals. The local Poisson model also ignores the mutation rate heterogeneity across patients. However, it uses a Poisson regression against the same set of covariates as the NIMBus model to compensate large-scale mutational heterogeneity across the genome.

### Global and local Binomial models

In [4], after pooling samples from a certain disease, a constant mutation rate was assumed at each single nucleotide over the genome. Hence, the number of mutations 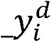 within a given region with length *l*_*i*_ follows a Binomial distribution as

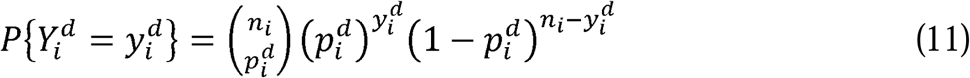

where 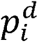 is the mutation rate at a single nucleotide. In a global Binomial model, 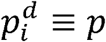 is assumed, and *p* is calculated in a genome-wide way. To remove the covariate effect, we may also assume a local Binomial model by using different 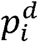 for different regions. Specifically, 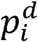 can be approximated by the length normalized 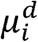 in NIMBus.

In order to check the degree of overdispersion in the mutation counts by Binomial assumption, we compared the observed and fitted mutation count data by Binomial distribution and provided the KS statistic in each cancer type. Specifically, we counted the number of mutations 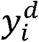 in 1mb bins generated in the Methods “Local background mutation rate estimation: (B)” section. Then the maximum likelihood estimate of mutation rate 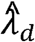 per position under the constant mutation rate assumption is calculated for cancer type *d*. Then we randomly generated simulated mutation counts 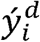 with 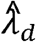 and calculated the KS statistic. We repeated the above process 100 times and plot the cumulative density function (C.D.F) of these KS statistics. A large KS statistic near 1 indicates larger overdispersion in the mutation count data. From Fig. S3, we showed that in all 7 cancer types, Binomial model provides poor fitting.

### Coding region annotation

We first extracted all the coding regions from the GENCODE v19 annotation. For annotation accuracy, we only selected the protein coding genes with gene_status labeled as “KNOWN” from the annotation. Then all the protein coding transcripts of the selected genes were selected. We merged multiple transcripts to get the final protein coding gene annotation as shown in Fig. S9. In total, 19,291 known protein-coding genes haven been used in this analysis.

## Noncoding annotations

We collected the full list of noncoding annotations to the best of our knowledge and customized it suitable for burden analysis. This list includes promoter regions, transcription start sites (TSS), translated regions (UTR), transcription factor binding sites (TFBS), enhancers, ultra-conserved, and ultra-sensitive sites. Promoters and TSS sites of known protein coding genes were defined as the 2500 and 100 nucleotides (nt) before the transcripts annotated by GENCODE v19. We also collected all the TFBS and enhancers from all tissues that are uniformly processed through the ENCODE pipeline. In addition, the ultra-conserved and ultra-sensitive sites were defined as those under positive selection during transcription regulations in our previous method FunSeq [41].

### Simulated variants for all cancer types

For each variant in a set of whole genome sequencing data, we tried to find a new position in a 100kb neighboring region (50k and 50k up and downstream each). Then we tested all the coding genes defined above on the original and simulated data set. Since the permuted size 100kb is relatively large as compared to the test region, a better method is supposed to give less or even no positives on the permuted data set. The Q-Q plots of P values of protein coding genes in both real and simulated data were given in Fig. S10.

### Mutation burden test for networks in the genome

In addition to testing single target regions, it is useful to extend our analysis to testing of networks. We took the KEGG pathway as a natural biological application of our network analysis[42]. Each coding gene represents a target region in the genome, and the gene set that makes up a pathway represents a network of genes. Since a KEGG pathway may consist of genes that are located on different chromosomes or regions of the genome, the mutation burden for a pathway will be heterogeneous. We assume that these heterogenous mutation burden levels are independent due to the disjoint, discontinuous association of each region. Therefore, for each pathway, we first determine the *p*-value of each coding gene in the pathway list using the local mutation burden calculations from NIMBus, and then combine them using Fisher’s method for a pathway associated *p*-value. This example can be seen in Figure 6.

**Figure 6.**
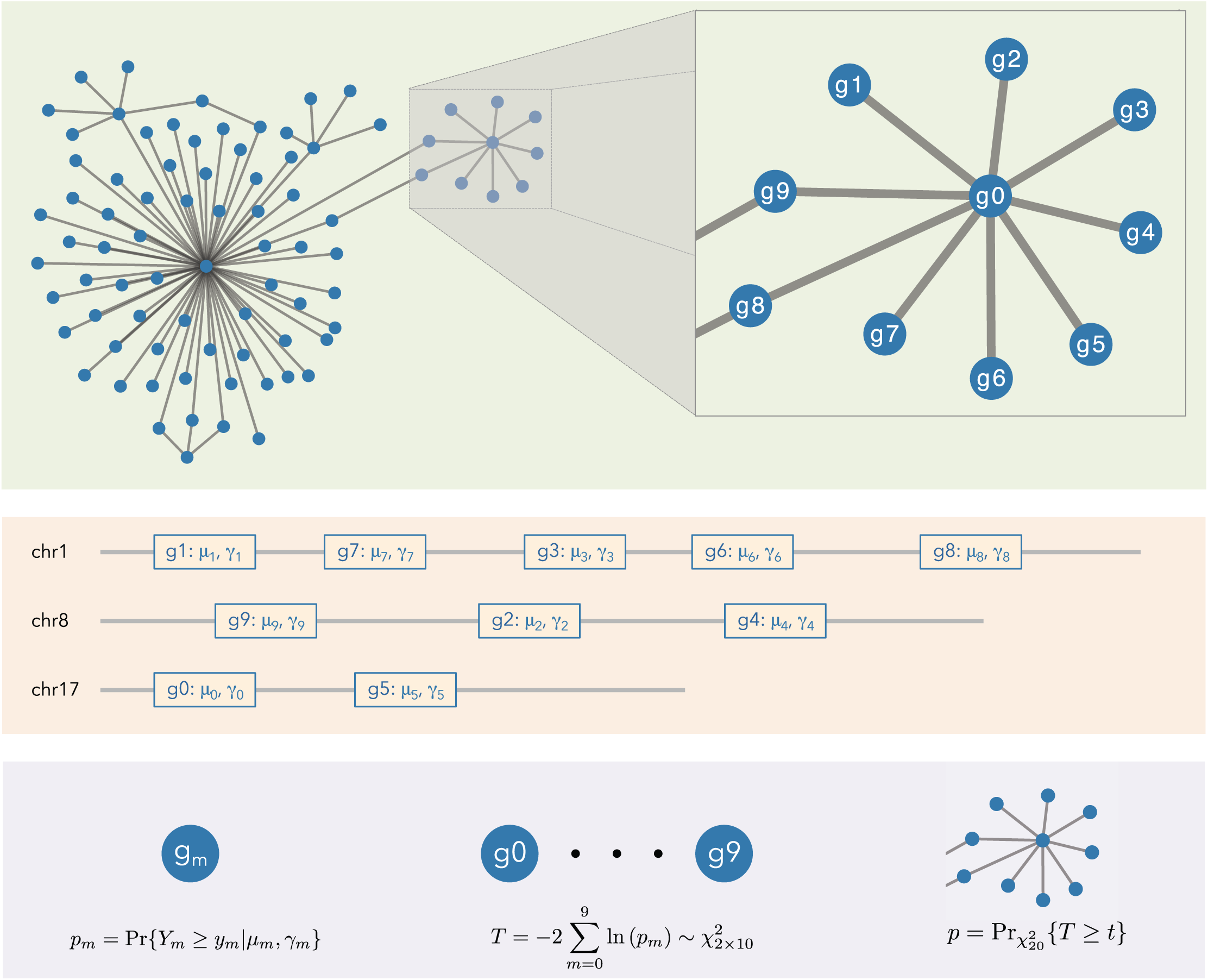
Schematic plot of network analysis: The associated values of *μ*_*m*_, *γ*_*m*_ are extracted from NIMBus for each of the *m* genes, potentially located on different chromosomes. A single p-value, *p*_*m*_, is obtained for each gene. Fisher’s method is used to combine all of the p-values into a final p-value for the network,*p*_*network*_.

In our analysis, given a network of regions consisting of *M* (*m* = 1,..,*M*) individual regions, each with*y*_*m*_ mutations, we can determine the *p*-value (*p*_*m*_) associated with each individual region based on *μ*_*m*_ and *γ*_*m*_, and then combine these *p*-values to produce a single *p*-value (*p*_*comb*_) associated with the network. To do this, we use Fisher’s method for combining *p*-values.

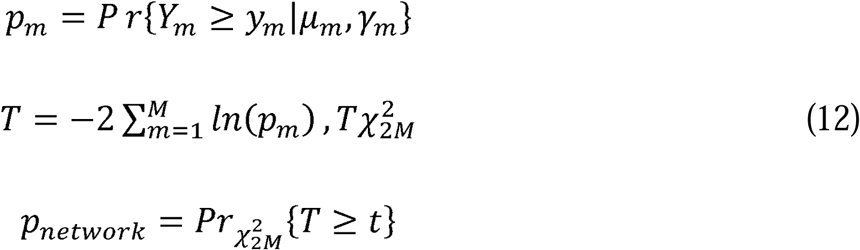

We took the KEGG pathway as a natural biological application of our network analysis. Each coding gene represents a target region in the genome, and the gene set that makes up a pathway represents a network of genes. Since a KEGG pathway may consist of genes that are located on different chromosomes or regions of the genome, the mutation burden for a pathway will be heterogeneous.

### Identification of predicted functional TF Motifs

Transcription Factor Binding Sites (TFBSs) that fall within DHS regions and also within a promoter region were identified in a cancer-type-specific manner. First, transcription start sites (TSSs) were identified upstream of both coding and non-coding transcripts using the GENCODE Hg19 annotation. Promoter regions were defined as the region of 1.5k BP upstream and 1k BP downstream of a TSS. Second, in order to map transcriptional activity, DHS signal tracks from ENCODE were intersected with these promoter regions in order to identify areas of likely functional significance. This mapping was completed in a cancer-cell-type-specific manner for a total of 14 cancer types with whole genome sequencing from PCAWG and corresponding cancer cell lines from ENCODE (Table S5). Third, TFBSs from ENCODE were identified that intersect with these promoter and DHS regions. As further screening step, RNA-seq expression data was used to limit TFs analyzed to those with non-negligible expression. 7 cancer types had associated RNA-seq data in ENCODE (Tier 1 annotation - Myeloid-MPN, Lung-SCC, Liver-HCC, Panc-AdenoCA, Breast-AdenoCa, Cervix-AdenoCA, Skin-Melanoma), 7 did not have matched RNA-seq data (Tier 2 annotation - Lymph-CLL, Lung-AdenoCA, Lymph-BNHL, ColoRect-AdenoCA, Myeloid-AML, Prost-AdenoCA, CNS-Medullo). For Tier 1 cancer types with RNA-seq data, the matched RNA-seq data, TFs analyzed were limited to those with average FPKM > 1 across replicates. 750 TFs were analyzed, 702 of which had RNA-seq-based expression level data available. For Tier 2 cancer types lacking this RNA-seq data, all 750 TFs were analyzed. The TFBSs identified were then aggregated at the gene level and carried forward to subsequent burdening analysis with NIMBus using the PCAWG variant call set.

### Benchmarking NIMBus

We first benchmarked our noncoding results by comparing them to those derived by using OncodriveFML (v. 2.1.3) [24]. OncodriveFML was run via command line in a Python 3.5 environment with the BBGLab bgparsers package (v. 0.7). In total, running our annotations and variants using OncodriveFML used 881.12 GB and took 43 hours. Significant results were identified using the Q value < 0.05.

Secondly, we benchmarked our results to the driver genes independently identified by the PCAWG Consortium [23]. Two cancers (Cervix-AdenoCA and Myeloid-AML) that were analyzed by NIMBus were not available for direct comparison in the PCAWG Consortium data (Text S2). Additionally, we used the well-known Cancer Gene Census COSMIC database to classify burdened genes according to their existing literature support. We downloaded the most current list from https://cancer.sanger.ac.uk/cosmic/download. More details can be found in Supplemental Text S2.

## Additional files

**Supplemental Figures 1-10**

**Supplemental Tables 1-5**

**Supplemental Texts 1-2**

The NIMBus model can flexibly be retrained and run using a new set of covariates, sequencing results, and target regions. Instructions and software are provided for this purpose at github.gersteinlab.org/nimbus.

## Abbreviations

BRCA: Breast cancer
GACA: gastric cancer
LICA: liver cancer
LUAD: Lung adenocarcinoma
PRAD: prostate cancer
MB: Medulloblastoma
PA: Pilocytic Astrocytoma

## Authors’ contributions

JZ, JL, PDM, and MG designed the study. JZ, JL, PDM, CY, DL, and MG wrote the manuscript. JZ, JL, PDM, CY, and LL wrote the framework and performed the noncoding analysis. JL and PDM worked on the network analysis and JL, PDM, and CY built the project website.

## Acknowledgements

This work was supported by the National Institutes of Health, AL Williams Professorship, and in part by the facilities and staff of the Yale University Faculty of Arts and Sciences High Performance Computing Center.

