## Supplement for "NIMBus: a Negative Binomial Regression based Integrative Method for Mutation Burden Analysis"

NIMBus Supplementary Document:

Table of Contents

Table S1	2

Figure S1	2

Figure S3	4

Figure S4	5

Figure S6	5

Table S2	6

Table S3	7

Figure S7	7

Figure S8	8

Figure S9	9

Figure S10	10

Table S1. Summary of WGS data

| **cancer** | **median** | **sd** |
| --- | --- | --- |
| BRCA | 3705 | 7300.526 |
| GACA | 14429.5 | 71372.080 |
| LICA | 8706.5 | 5522.917 |
| LUAD | 21287 | 35610.839 |
| MB | 965 | 1196.036 |
| PA | 70 | 114.414 |
| PRAD | 4927 | 2764.873 |

Figure S1. Pie chart of sample numbers of the WGS data

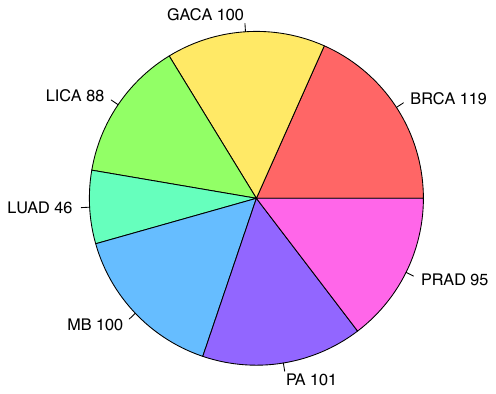

Among these samples, 100 stomach cancer samples were from Wang *et al* [1] and 95 prostate cancer samples were obtained from our collaborators. The remaining comes from samples published by Alexandrov *et al* [2].

Figure S2. Schematic sketch of optimal and approximate local mutation rate estimation

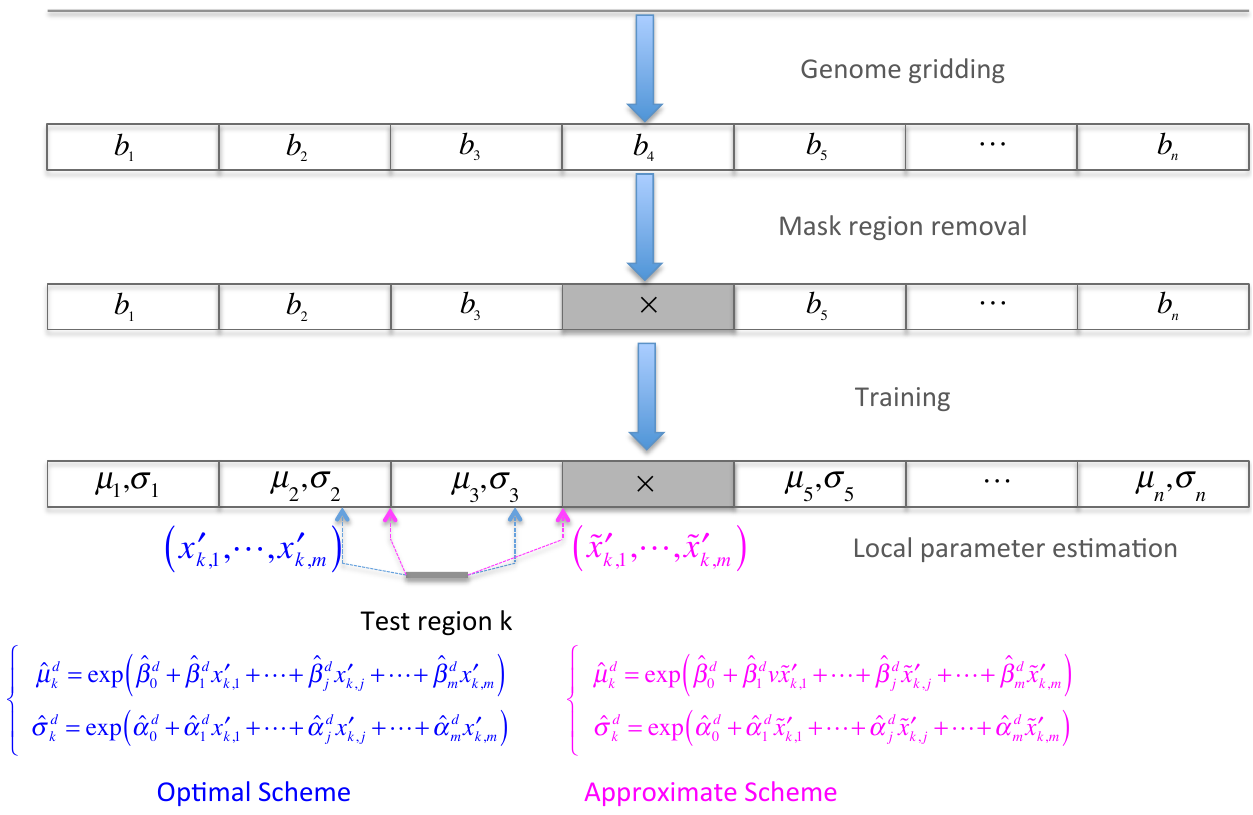

Text S1. Overview of approximation scheme

In real data analysis there are usually millions of regions to be tested and for each region it needs to process 381 features. Hence, the above optimal scheme is usually computational expensive. Here we proposed an approximation scheme to calculate $\hat{\mu}_{k}^{d}$ and $\hat{\sigma}_{k}^{d}$. Instead of using covariates for the extended bin centered at target region $k$, we used the values for the nearest training bin $\left( \tilde{x}_{k,1}^{'},\cdots,\tilde{x}_{k,m}^{'} \right)$ (magenta parts in Fig. S2), and burden tests are performed after length adjustment. Since $\left( \tilde{x}_{k,1}^{'},\cdots,\tilde{x}_{k,m}^{'} \right)$ has already been pre-calculated during the training process, our approximation scheme significantly reduced the computation burden for tests.

Figure S3. KS statistic of the observed and fitted mutation counts using Binomial distribution

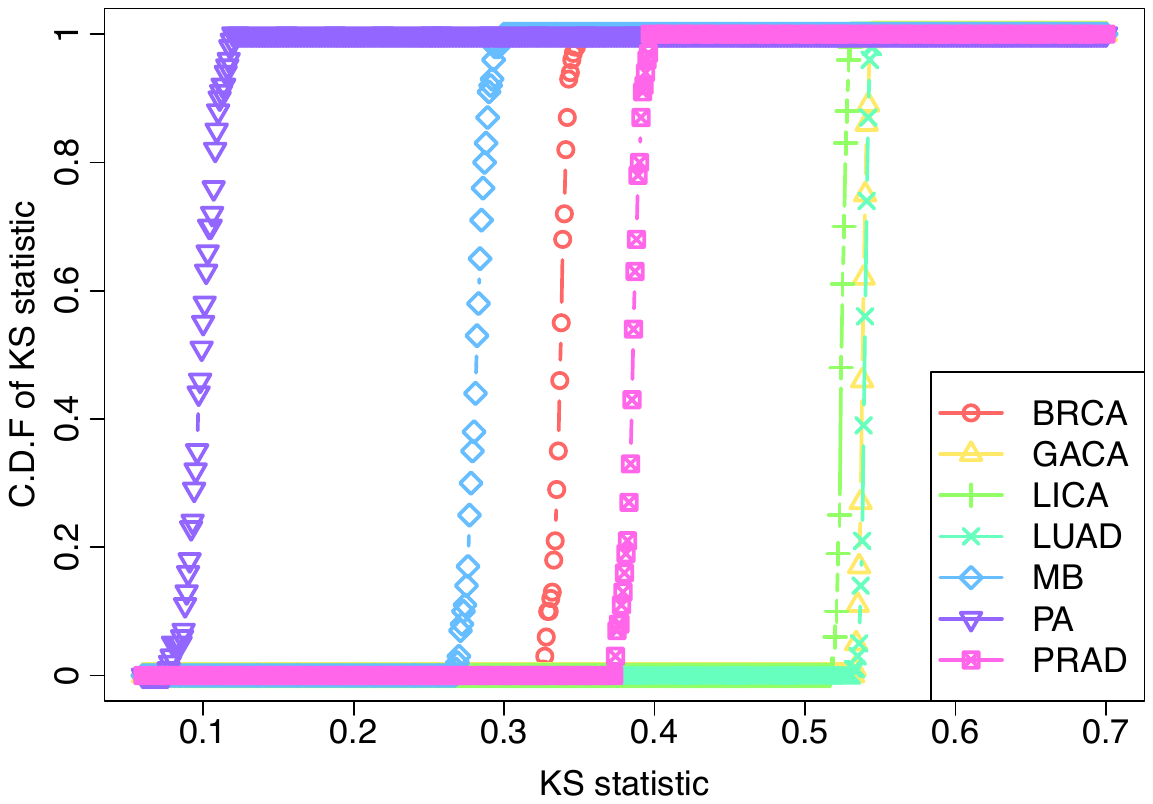

Figure S4. Mutation rates are severely affected by many genomic features in breast cancer in the first 70 1mb bins on chromosome 1. Mutation counts (black line and left y axis) and other genomic features (red line and right y axis) are normalized in a genome-wide way.

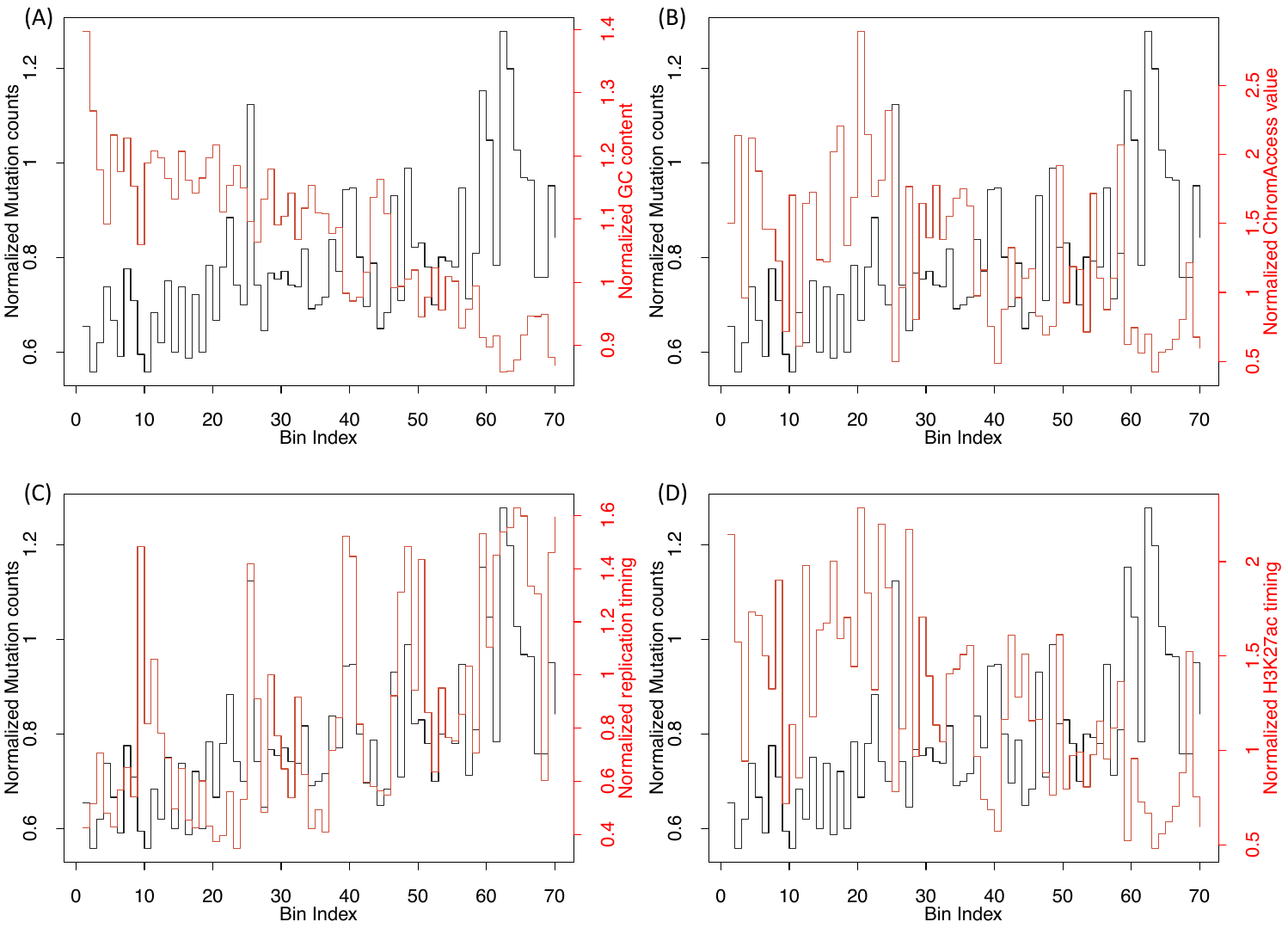

Figure S6. (A) Features in breast and brain related tissues are correlated; (B) scatter plot of observed and predicted mutations by regression for all four models. Red line represents the diagonal line and Pearson correlations are also given.

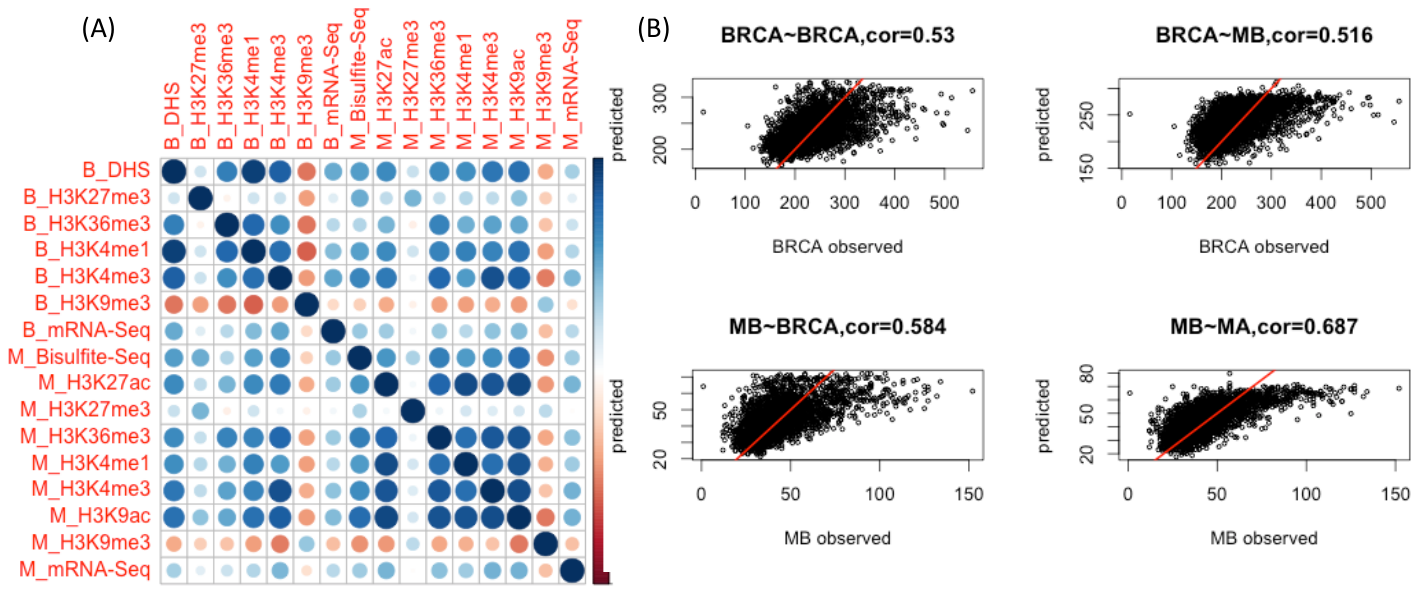

Table S2. Four regression models using matched and unmatched genomic features

| _Mutations_ ^Covariates^ | BRCA | MB |
| --- | --- | --- |
| BRCA | 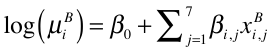 | 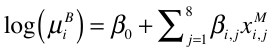 |
| MB | 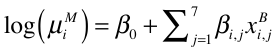 | 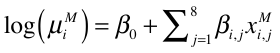 |

Table S3. Relative error for models using matched and unmatched genomic features

| _Mutations_ ^Covariates^ | BRCA | MB |
| --- | --- | --- |
| BRCA | 0.128 | 0.135 |
| MB | 0.195 | 0.183 |

Figure S7. (A) Cumulative proportion of variance explained by the number of PCs; (B) Boxplot of Pearson correlations of top PCs to mutation counts data in different cancer types.

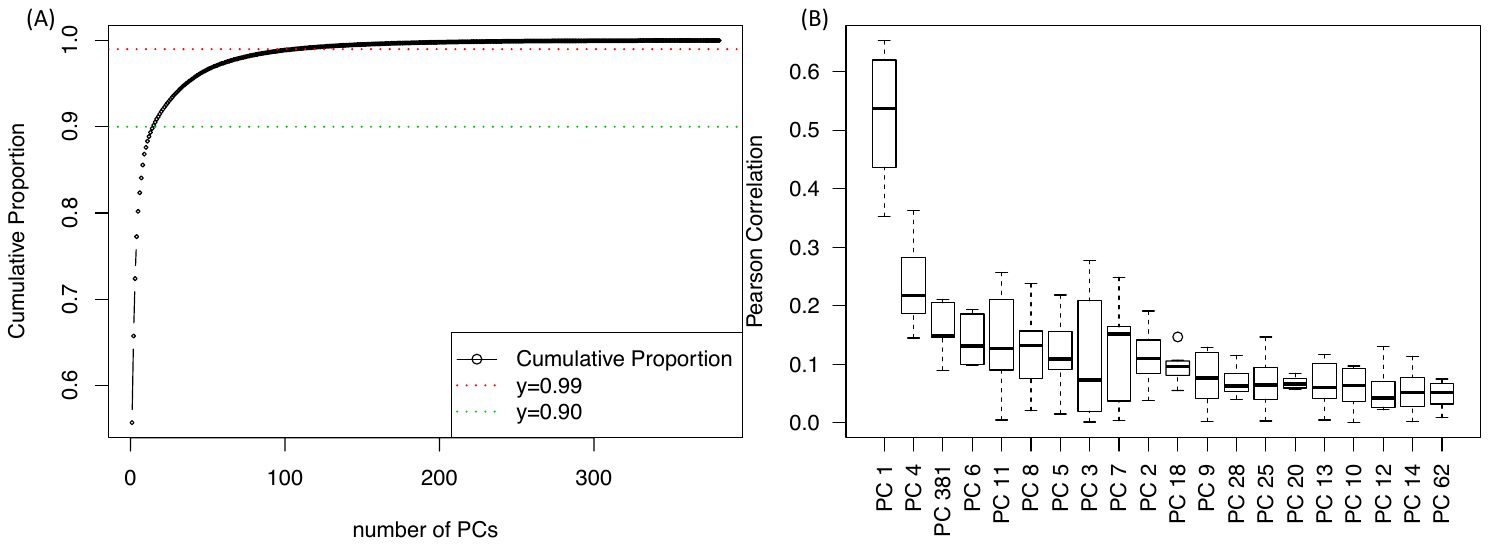

Figure S8. Scatter plot of observed and predicted mutation rate. Red line represents the diagonal line.

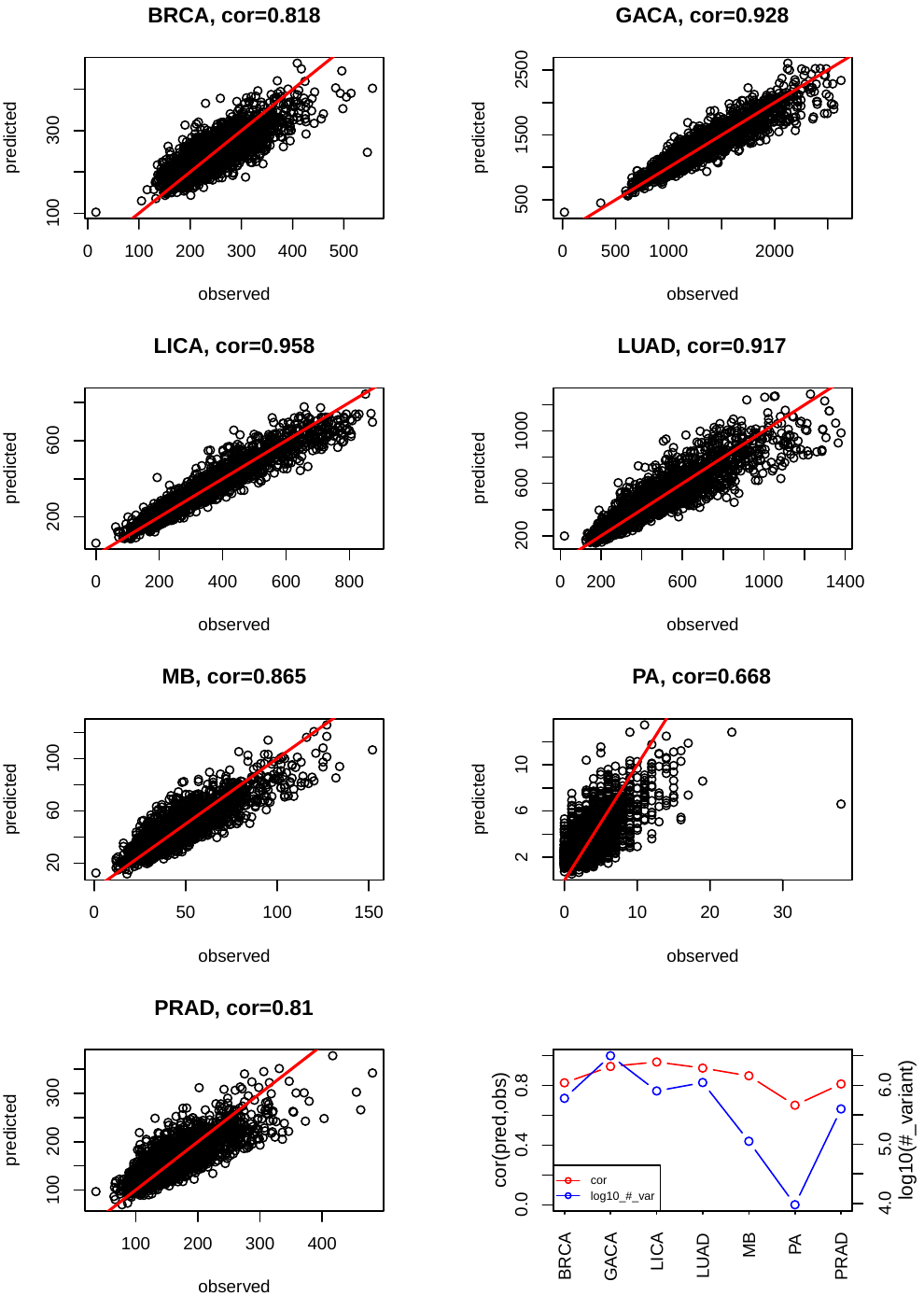

Figure S9. Example of protein coding gene merging

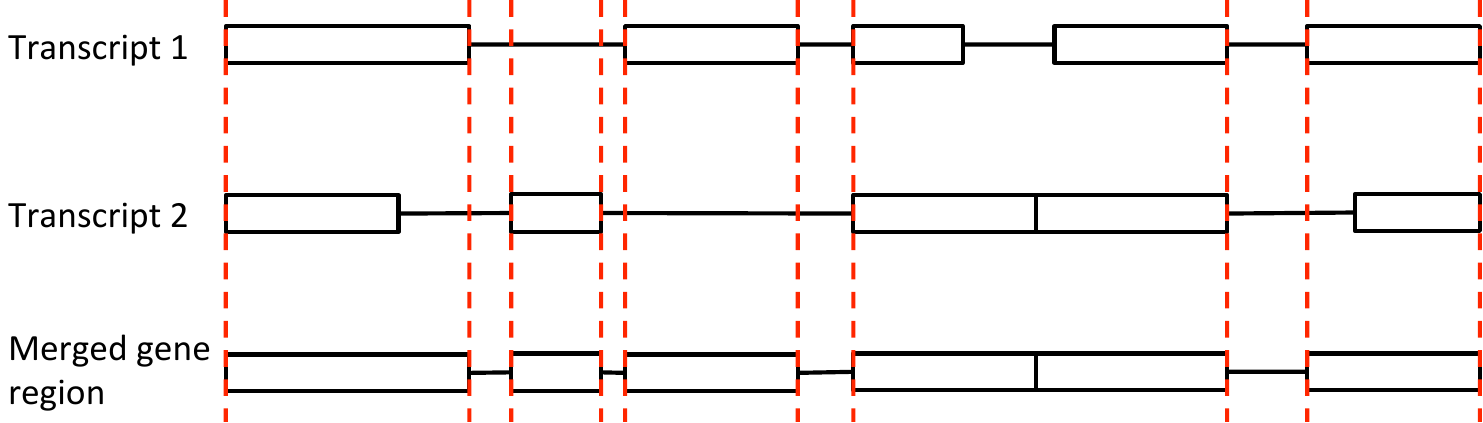

Table S4 significant genes after P value combination

| **Rank** | **Gene** | **Adjust P** | **PubMed ID** |
| --- | --- | --- | --- |
| 1 | TP53 | 4.33E-139 | 17401424 |
| 2 | DDX3X | 3.65E-18 | 22820256 |
| 3 | KRAS | 2.56097E-06 | 19847166 |
| 4 | MUC4 | 4.46636E-06 | 19935676 |
| 5 | CDH1 | 3.06519E-05 | 10973239 |
| 6 | ARID1A | 0.000236255 | 22037554 |
| 7 | SMARCA4 | 0.000377726 | 18386774 |
| 8 | FGFR1 | 0.000742868 | 23817572 |
| 9 | OTOP1 | 0.003694811 | NA |
| 10 | SPOP | 0.004493795 | 22610119 |
| 11 | STK11 | 0.004493795 | 15021901 |
| 12 | PTEN | 0.004812088 | 9697695 |
| 13 | SMO | 0.012684045 | 9422511 |
| 14 | TAS2R31 | 0.033983962 | NA |
| 15 | TBC1D29 | 0.034276895 | NA |

Figure S10. Q-Q plots of gene regions on real and simulated data

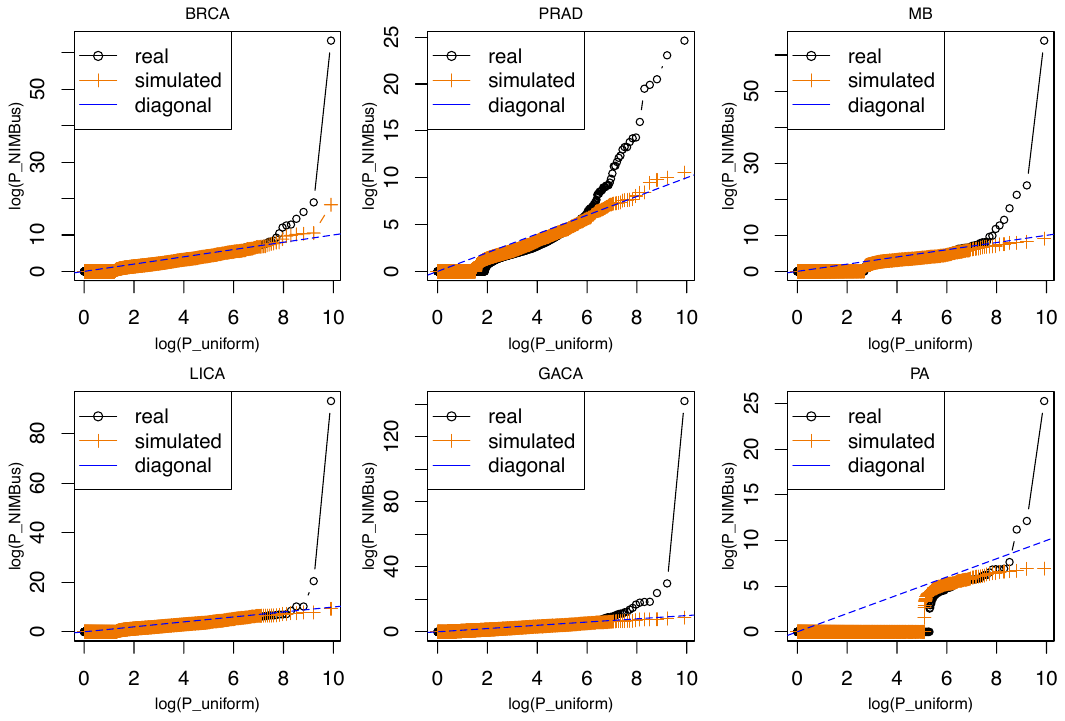

Table S5. Cancer type and assay matching

| **ENCODE DNase** | **PCAWG cancer type** |
| --- | --- |
| SK-MEL-5_ENCSR000FEK_rep1 | Skin-Melanoma |
| SK-MEL-5_ENCSR000FEK_rep2 | Skin-Melanoma |
| HeLa-S3_ENCSR000ENO_rep1 | Cervix-AdenoCA |
| HeLa-S3_ENCSR000ENO_rep2 | Cervix-AdenoCA |
| HeLa-S3_ENCSR000EJT_rep1 | Cervix-AdenoCA |
| MCF-7_ENCSR000EPH_rep1 | Breast-AdenoCa |
| MCF-7_ENCSR000EPH_rep2 | Breast-AdenoCa |
| MCF-7_ENCSR000EPJ_rep1 | Breast-AdenoCa |
| MCF-7_ENCSR000EPJ_rep2 | Breast-AdenoCa |
| MCF-7_ENCSR000EKX_rep1 | Breast-AdenoCa |
| MCF-7_ENCSR000EKW_rep1 | Breast-AdenoCa |
| MCF-7_ENCSR000EKV_rep1 | Breast-AdenoCa |
| MCF-7_ENCSR000EKZ_rep1 | Breast-AdenoCa |
| T47D_ENCSR000ELT_rep1 | Breast-AdenoCa |
| PC-9_ENCSR940NLN_rep1 | Lung-AdenoCA |
| PC-9_ENCSR940NLN_rep2 | Lung-AdenoCA |
| Karpas-422_ENCSR019JDO_rep2 | Lymph-BNHL |
| Karpas-422_ENCSR019JDO_rep1 | Lymph-BNHL |
| CMK_ENCSR000EMN_rep1 | Myeloid-AML |
| Caco-2_ENCSR000EMI_rep1 | ColoRect-AdenoCA |
| Caco-2_ENCSR000EMI_rep2 | ColoRect-AdenoCA |
| K562_ENCSR000EKP_rep1 | Myeloid-AML |
| K562_ENCSR000EKQ_rep1 | Myeloid-AML |
| K562_ENCSR000EKS_rep1 | Myeloid-AML |
| K562_ENCSR000EPC_rep1 | Myeloid-AML |
| K562_ENCSR000EPC_rep2 | Myeloid-AML |
| K562_ENCSR921NMD_rep1 | Myeloid-AML |
| K562_ENCSR921NMD_rep2 | Myeloid-AML |
| Panc1_ENCSR000EPT_rep1 | Panc-AdenoCA |
| Panc1_ENCSR000EPT_rep2 | Panc-AdenoCA |
| HCT116_ENCSR000ENM_rep2 | ColoRect-AdenoCA |
| HCT116_ENCSR000ENM_rep1 | ColoRect-AdenoCA |
| WERI-Rb-1_ENCSR000EQL_rep1 | ### |
| WERI-Rb-1_ENCSR000EQL_rep2 | ### |
| OCI-LY7_ENCSR489NAM_rep1 | Lymph-BNHL |
| OCI-LY7_ENCSR489NAM_rep2 | Lymph-BNHL |
| HepG2_ENCSR000EJV_rep1 | Liver-HCC |
| HepG2_ENCSR000ENP_rep1 | Liver-HCC |
| HepG2_ENCSR000ENP_rep2 | Liver-HCC |
| HAP-1_ENCSR620QNS_rep1 | Myeloid-MPN |
| HAP-1_ENCSR620QNS_rep2 | Myeloid-MPN |
| A549_ENCSR000EIE_rep1 | Lung-SCC |
| A549_ENCSR000ELW_rep1 | Lung-SCC |
| A549_ENCSR000ELW_rep2 | Lung-SCC |
| NT2-D1_ENCSR000EPS_rep2 | ### |
| NT2-D1_ENCSR000EPS_rep1 | ### |
| SJCRH30_ENCSR700ILE_rep1 | ### |
| BE2C_ENCSR000EMD_rep1 | ### |
| BE2C_ENCSR000EMD_rep2 | ### |
| HL-60_ENCSR000ENU_rep2 | Myeloid-AML |
| HL-60_ENCSR000ENU_rep1 | Myeloid-AML |
| SK-N-MC_ENCSR000EPY_rep1 | ### |
| SK-N-MC_ENCSR000EPY_rep2 | ### |
| LNCaP-clone-FGC_ENCSR000EKT_rep1 | Prost-AdenoCA |
| LNCaP-clone-FGC_ENCSR000EPF_rep1 | Prost-AdenoCA |
| LNCaP-clone-FGC_ENCSR000EPF_rep2 | Prost-AdenoCA |
| Daoy_ENCSR346IHH_rep2 | CNS-Medullo |
| Daoy_ENCSR346IHH_rep1 | CNS-Medullo |
| KBM-7_ENCSR426IEA_rep2 | Myeloid-MPN |
| KBM-7_ENCSR426IEA_rep1 | Myeloid-MPN |

Text S2: Benchmarking Methods Details

First, the intersection was found between burdened noncoding regions identified by NIMBus and identified by using OncodriveFML. Secondly, the intersection was found between burdened noncoding regions identified by NIMBus and the PCAWG Consortium [3]. The CDS regions from the PCAWG data were not used for this comparison. Furthermore, two cancers analyzed by NIMBus (Cervix-AdenoCA and Myeloid-AML) were not available for direct comparison in the PCAWG data. Meta-Adenocarcinoma and Meta-Myeloid were available in the PCAWG data, but are broader in scope than the cancer types used in NIMBus. Thus, Cervix-AdenoCA and Myeloid-AML were excluded from this comparison.

Lastly, the shared burdened regions and the regions unique to each method (NIMBus, OncodriveFML, and the PCAWG Consortium) were compared to the Tier 1 and Tier 2 cancer genes as identified by the COSMIC Cancer Gene Census [4]. Tier 1 genes possess strong experimental support as being related to cancer whereas Tier 2 genes possess developing experimental evidence.
